# Short-term high fat diet feeding of mice suppresses catecholamine-stimulated Ca^2+^ signalling in hepatocytes and intact liver

**DOI:** 10.1101/2022.08.28.505514

**Authors:** Robert P. Brumer, Juliana C. Corrêa-Velloso, Samantha J. Thomas, Oleta A. Sandiford, Andrew P. Thomas, Paula J. Bartlett

## Abstract

Excess consumption of carbohydrates, fat, and calories leads to non-alcoholic fatty liver disease (NAFLD) and hepatic insulin resistance; major factors in the pathogenesis of type II diabetes. Hormones and catecholamines acting through G-protein coupled receptors (GPCRs) linked to phospholipase C (PLC) and increases in cytosolic Ca^2+^ ([Ca^2+^]_c_) regulate many metabolic functions of the liver. In the intact liver, catabolic hormones such as glucagon, catecholamines and vasopressin integrate and synergize to regulate the frequency and extent to which [Ca^2+^]_c_ waves propagate across hepatic lobules to control metabolism. Dysregulation of hepatic Ca^2+^ homeostasis has been implicated in the development of metabolic disease, but changes in hepatic GPCR-dependent Ca^2+^ signalling have been largely unexplored in this context. We show that short-term, 1-week, high fat diet (HFD) feeding of mice attenuates norepinephrine-stimulated Ca^2+^ signalling, reducing the number of cells responding and suppressing the frequency of [Ca^2+^]_c_ oscillations in both isolated hepatocytes and intact liver. The 1-week HFD feeding paradigm did not change basal Ca^2+^ homeostasis; endoplasmic reticulum Ca^2+^ load, store-operated Ca^2+^ entry and plasma membrane Ca^2+^ pump activity were unchanged compared to low fat diet (LFD) fed controls. However, norepinephrine-induced IP_3_ production was significantly reduced after HFD feeding, demonstrating an effect of HFD on receptor-stimulated PLC activity. Thus, we have identified a lesion in the PLC signalling pathway induced by short-term HFD feeding, which interferes with hormonal Ca^2+^ signalling in isolated hepatocytes and the intact liver. These early events may drive adaptive changes in signalling, which lead to pathological consequences in fatty liver disease.

**Key points summary:**

- Non-alcoholic fatty liver disease (NAFLD) is a growing epidemic.
- In healthy liver, the counteracting effects of catabolic and anabolic hormones regulate metabolism and energy storage as fat. Hormones and catecholamines promote catabolic metabolism via increases in cytosolic Ca^2+^ ([Ca^2+^]_c_).
- We show that 1 week high fat diet (HFD) feeding of mice attenuated the Ca^2+^ signals induced by physiological concentrations of norepinephrine. Specifically, HFD suppressed the normal pattern of periodic [Ca^2+^]_c_ oscillations in isolated hepatocytes and disrupted the propagation of intralobular [Ca^2+^]_c_ waves in the intact perfused liver.
- Short-term HFD inhibited norepinephrine-induced inositol 1,4,5-trisphosphate (IP_3_) generation, but did not change basal endoplasmic reticulum Ca^2+^ load or plasma membrane Ca^2+^ fluxes.
- We propose that impaired Ca^2+^ signalling plays a key role in the earliest phases of the etiology of NAFLD, and is responsible for many of the ensuing metabolic and related dysfunctional outcomes at the cellular and whole tissue level.

## Introduction

Non Alcoholic Fatty liver disease (NALFD) is a global health concern and is currently the most common liver disease worldwide (Fazel *et al*., 2016; Muzurović *et al*., 2021). NAFLD is characterized by the abnormal storage of fat in the liver and arises due to caloric oversupply from excess carbohydrate and fat (Fung *et al*., 2001). It is highly prevalent, affecting 20-30% of Americans (Sayiner *et al*., 2016), and has a strong epidemiological link with type 2 diabetes mellitus, affecting about 60% of diabetics (Dai *et al*., 2017). End-stage NAFLD is an inflammatory disease called non-alcoholic steatohepatitis (NASH), which causes cirrhosis (scarring of the liver) and can progress to hepatocellular carcinoma, and is currently the second highest indication for liver transplant in the US (Wong *et al*., 2015). Current treatment options are limited and do not provide a cure. Identification of novel drug targets for the effective treatment of liver disease requires a detailed understanding of the cellular and molecular mechanisms promoting NAFLD and NASH. Hepatic lipid and glucose metabolism are maintained by the interplay of counteracting hormones, intracellular signalling pathways and transcription factors, allowing for regulation of hepatic metabolism and the maintenance of euglycemia (Vollmers *et al*., 2009; Rui, 2014; Miller & Birnbaum, 2016). In healthy individuals the counteracting effects of insulin and glucagon, in concert with other glucose producing (glucogenic) hormones, maintain the balance of storage and output of energy metabolites by the liver. A role of insulin resistance in the manifestation of metabolic syndrome is well established, but changes in the counteracting catabolic signals have been largely unexplored.

Hormones and catecholamines acting through receptors coupled to phospholipase C (PLC) activation regulate many metabolic functions of the liver by stimulating the formation of inositol 1,4,5-trisphosphate (IP_3_) and diacylglycerol (DAG), leading to increases in cytosolic [Ca^2+^]_c_ (Rooney *et al*., 1989; Thomas *et al*., 1995; Bartlett *et al*., 2014). It is now well established that glucagon also regulates gluconeogenesis through a Ca^2+^-dependent mechanism, via cAMP-dependent phosphorylation of IP_3_ receptors (Hajnóczky *et al*., 1993; Rooney *et al*., 1996; Ozcan *et al*., 2012; Wang *et al*., 2012; Perry *et al*., 2020). Thus, at physiological levels glucagon does not act alone and requires the presence of PLC-linked hormones to maintain euglycemia. Ca^2+^ responses elicited by glucogenic hormones also regulate hepatic lipid metabolism, suppressing fatty acid synthesis and promoting β-oxidation. Hormone-induced Ca^2+^ signals in hepatocytes manifest as baseline seperated [Ca^2+^]_c_ oscillations that propagate through the cytoplasm as intracelluar [Ca^2+^]_c_ waves (Rooney *et al*., 1989; Thomas *et al*., 1991; Hajnóczky & Thomas, 1997; Gaspers *et al*., 2014). Moreover, in the intact liver these intracellular oscillatory [Ca^2+^]_c_ waves pass through gap junctions to generate intercelluar [Ca^2+^]_c_ waves (Robb-Gaspers & Thomas, 1995; Gaspers & Thomas, 2005; Gaspers *et al*., 2019). Hormone dose-dependence is encoded in the oscillation frequency and the radial spread of intercellular [Ca^2+^]_c_ waves in the liver, which both increase with increasing agonist dose. We have shown that in the intact liver, glucogenic PLC-linked hormones such as catecholamines and vasopressin integrate with glucagon and regulate the frequency and extent to which [Ca^2+^]_c_ waves propagate across the hepatic lobules to control hepatic metabolism (Gaspers & Thomas, 2005; Bartlett *et al*., 2014; Gaspers *et al*., 2019). Importantly, there is zonal segregation of metabolic functions in the liver, with gluconeogenesis occurring predominantly in the periportal zone of the hepatic lobule and lipogenesis occurring predominantly in the pericentral zone. Significantly in this context, hormone- and catecholamine-induced Ca^2+^ signals are initiated in the periportal hepatocytes of liver lobules and must propagate across the lobule into the pericentral zone to fully integrate the regulation of hepatic metabolism in the intact organ.

Mouse models of obesity have revealed that fatty liver is associated with dysregulated hepatic Ca^2+^ homeostasis. A reduction in SERCA activity has been found in the ob/ob leptin-deficient mouse, and this is associated with decreased endoplasmic reticulum (ER) Ca^2+^ load and increases in markers of ER stress (Fu *et al*., 2011; Arruda *et al*., 2017). In addition, increased transfer of Ca^2+^ from ER to mitochondria leading to mitochondrial Ca^2+^ overload and reactive oxygen species (ROS) production has been reported (Arruda *et al*., 2014; Feriod *et al*., 2017). A reduction in store-operated Ca^2+^ entry (SOCE) has also been indicated as a factor in the dysregulation of glucose metabolism and development of hepatic steatosis (Wilson *et al*., 2015; Arruda *et al*., 2017). However, past studies in animal models have relied on germline endocrine defects in genetic models, or long-term dietary feeding. By contrast, our mouse study has identified a Ca^2+^ signalling defect after short-term exposure to high fat diet (HFD). Specifically, we have found a dramatic attenuation of PLC-linked [Ca^2+^]_c_ signalling at the cellular and whole organ level following 1 week of HFD. This suppression of hormone-stimulated Ca^2+^ signalling is caused by a decrease in IP_3_ production, with no change in IP_3_ receptor (IP_3_R) function, ER Ca^2+^ sequestration or SOCE. We demonstrate that suppression of norepinephrine-dependent [Ca^2+^]_c_ signalling by HFD is an early event in the development of hepatic steatosis, preceding other reported changes in Ca^2+^ homeostasis (Fu *et al*., 2011; Arruda *et al*., 2014; Feriod *et al*., 2017). The consequence of this reduced glucogenic signalling capacity is predicted to disrupt the balance between catabolic and anabolic signalling in the liver, giving rise to metabolic dysfunction and steatosis.

## Methods

Mice used for experiments were either purchased from Taconic Biosciences (C57BL/6N) or sourced from in-house breeding and were housed in an AAALAC accredited facility on the Rutgers NJMS campus. All procedures involving mice were approved by the Institutional Animal Care and Use Committee. Mice were fed LFD or HFD from age 5-6 weeks until the experimental endpoint. The diets used were LFD 5755 and HFD 58V8 from TestDiet®, which contain 10% and 45% calories from fat in the LFD and HFD, respectively. Before surgical procedures mice were anesthetized with 80 mg/kg ketamine and 12 mg/kg xylazine, or 60 mg/kg pentobarbital sodium, livers were harvested for tissue, or perfused to isolate hepatocytes or intravital imaging of the intact liver, exsanguination occurs during the procedure and whilst the mouse is anesthetized.

### Generation of liver specific-GCaMP6f mouse model

GCaMP6f floxed mice, B6J.Cg-Gt(ROSA)26Sor^tm95.1(CAG-GCaMP6f)Hze^/MwarJ, (The Jackson Laboratory; 028865) were bred with Albumin-Cre mice, B6N.Cg-Speer6-ps1^Tg(Alb-cre)21Mgn^/J (The Jackson Laboratory; 018961). The GCaMP6f floxed mice are a B6/J-derived mouse line that contain the Nnt^C57BL/6J^ mutation in the nicotinamide nucleotide transhydrogenase (Nnt) gene, which catalyzes the reduction of NADP^+^ to NADPH in the mitochondria. Since this mutation affects normal mitochondrial metabolism, we crossed the heterozygous offspring and selected mice homozygous for GCaMP6f floxed, Albumin-cre and wild type Nnt to establish our Liver specific-GCaMP6f mouse line; GCaMP6f^hep/cre^. Genotyping was performed by Transnetyx^®^ using primers validated by The Jackson Laboratory.

### Hepatocyte isolation and culture

Hepatocytes were prepared by collagenase perfusion of the mouse liver *in situ* using isolation buffer composed of 137 mM NaCl, 5.4 mM KCl, 0.64 mM NaH_2_PO_4_, 0.85 mM Na_2_HPO_4_, 23.8 mM NaHCO_3_, 5 mM glucose, 10 mM HEPES gassed with 5% CO_2_/95% O_2_ at pH7.4. An intravenous catheter was inserted into the inferior vena cava and the liver perfused with Ca^2+^-free isolation buffer containing 0.5 mM EGTA. The portal vein was immediately severed to enable buffer to flow retrograde through the liver (3 ml/min) and the perfusion continued for 10 min. This was followed by 5-10 min perfusion with collagenase-containing isolation buffer without EGTA and with 2.5 mM CaCl_2_, 0.2% w/v dialyzed bovine serum albumin (BSA) fraction V (Sigma A7906) and 0.5 mg/ml type IV collagenase from *Clostridium histolyticum* (Sigma-Aldrich C5138). Perfusion buffers were heated to 37 °C and equilibrated with 5% CO_2_/95% O2. Once digested, the liver was removed and hepatocytes were mechanically dissociated in isolation buffer containing 2.5 mM CaCl_2_ at 0 °C, followed by filtration through a cell strainer. Cells were centrifuged at 50 g for 2 min at 4 °C, washed three times in the isolation buffer and maintained on ice prior to plating.

For experiments using freshly-isolated hepatocytes, cells were cultured in Williams E medium containing 5 units ml/L penicillin 5 µg/mlstreptomycin, 50 µg/L gentamicin, and 2 mM L-glutamine in a humidified incubator at 37 °C maintained at 5% CO_2_. For overnight culture, cells were plated in the same media with the addition of 5% fetal bovine serum (FBS) and 140 nM insulin for 2 hours, then switched to 1% FBS and 250 pM insulin. Where possible experiments were performed in freshly isolated hepatocytes within 1 hour of plating.

### Intact perfused liver

For the *ex vivo* perfused liver preparation, a metal cannula was inserted into the portal vein and secured tightly with suture thread. The liver perfusion buffer was composed of 137 mM NaCl, 4.2 mM NaHCO_3_, 5.4 mM KCl, 0.81 mM MgSO_4_, 0.4 mM NaH_2_PO_4_, 0.44 mM KH_2_PO_4_, 1.3 mM CaCl_2_, 20 mM HEPES (pH 7.4), 5 mM glucose, 0.1X amino acids (Gibco; 11140076); room temperature. After portal vein cannulation, the inferior vena cava was cut to allow perfusion buffer outflow. The liver was then carefully excised from the animal and transferred to a confocal microscope stage, where intravital imaging was performed during continuous perfusion at 3 ml/min. Hormones were added to the perfusate as required. GCaMP6f^hep/cre^ mice bred in-house were fed with the required experimental diet for 1 week prior to experimentation.

### Single Cell Calcium Measurements

Hepatocytes were isolated by collagenase perfusion of livers from LFD or HFD as described above (Rooney *et al*., 1989). Cells were plated onto glass coverslips coated with collagen (type I rat tail, Sigma; 10 μg/cm^2^) and maintained in a humidified atmosphere of 5% CO_2_ and 95% air at 37 °C in Williams’ Medium E supplemented with penicillin, (10 units/ml), streptomycin (10 μg/ml), gentamycin sulfate (50 μg/ml), glutamine (2 mM), and insulin (140 nM). Adhered hepatocytes were loaded with 5 μM fura-2/AM and 0.02% (w/v) Pluronic F-127 for 20 min in HBSS composed of 25 mM HEPES, 121 mM NaCl, 5 mM NaHCO_3_, 4.7 mM KCl, 1.2 mM KH_2_PO_4_, 1.2 mM MgSO_4_, 1.3 mM CaCl_2_, 5 mM glucose, 0.04 mM probenecid, and 0.25% (w/v) fatty acid-free BSA, pH 7.4. After washing, the cells were transferred to a thermostatically controlled microscope imaging chamber and maintained at 37 °C on the microscope stage. Fluorescence images (excitation, 340 and 380 nm; emission, 420–600 nm) were acquired every 1-3s using a cooled charged-coupled device (CCD) camera as previously described (Rooney *et al*., 1989).

### Photorelease of caged IP_3_

Isolated hepatocytes were transfected with GCaMP6f (Chen *et al*., 2013) cDNA using nucleofection (Lonza) and cultured overnight in Williams E medium. Hepatocytes were then incubated in HBSS and loaded with the membrane permeant form of caged IP_3_ (2 µM, D-2, 3-O-isopropylidene6-O-(2-nitro-4,5-dimethoxy)benzyl-myo-inositol1,4,5-trisphosphate-hexakis(propionoxymethyl) ester (Li *et al*., 1998); Sichem GmbH) for 1 h at room temperature. After washing, the cells were transferred to a thermostatically-controlled microscope imaging chamber and maintained at 37 °C on the microscope stage. GCaMP6f fluorescence images (excitation 488 nm; emission 510 nm long pass) were acquired at 2 Hz. After recording an initial baseline period, continuous photorelease of caged IP_3_ was achieved with 50 ms exposure to 380 nm light at 2 Hz interlaced with the sequential GCaMP6f image acquisition (Bartlett *et al*., 2020).

### Permeabilized cell IP_3_ receptor function assay

The method was adapted from our previous work using compartmentalized fura-2 FF to measure changes in ER Ca^2+^ in response to IP_3_ (Hajnóczky *et al*., 1994; Hajnóczky & Thomas, 1997; Gaspers *et al*., 2021), except that ER Ca^2+^ was monitored using a plate reader. Freshly isolated hepatocytes were plated into a 96 well plate (30,000 cells/well) for 1 h, prior to loading with 5 μM fura-2 FF/AM for 1 h in HBSS (as above), to load the dye into intracellular compartments as well as the cytosol. Hepatocytes were then permeabilized with 25 µg/ml digitonin for 10 min in intracellular medium (ICM) composed of 120 mM KCl, 10 mM NaCl, 1.2 mM KH_2_PO_4_, 2 mM MgCl_2_, 20 mM Tris-HEPES, 1 mM EGTA, 0.5 mM CaCl_2_, 0.1X Protease Inhibitor (Calbiochem), 1μg/ml pepstatin, 1μg/ml oligomycin, 5 μM rotenone at pH 7.2, followed by washing 3 times with ICM to remove the released cytosolic fura-2 FF. The digitonin-permeabilization procedure leaves the ER-compartmentalized fura-2 FF intact and allows direct access to the IP_3_R by added IP_3_. Changes in ER Ca^2+^ were monitored with excitation at 340 nM and 380 nM (bandwidth 9 nM) and emission 510 nM (bandwidth 15 nM; cutoff 495 nM) at 4 s intervals using a plate reader (FlexstationIII, Molecular Devices) with micro fluidic addition capabilities during continuous data acquisition. Ca^2+^ uptake into the ER was initiated with 2 mM ATP to allow the stores to load with Ca^2+^ prior to addition of a range of IP_3_ concentrations (10 nM-10 μM), a maximum dose of IP_3_ (10 μM), followed by 10 μM ionomycin.

### Tissue histology

Fresh frozen liver was prepared by placing a piece of tissue in optimal cutting temperature (OCT) compound and flash freezing using solid isopentane on liquid nitrogen. Liver tissues were sectioned at 7-10 μm and fixed with 60% ethanol or 10% formalin and rinsed in 60% ethanol. For Oil Red O staining, a saturated solution (1% Oil Red O in 99% isopropyl alcohol) was diluted 3:2 in H2O and the liver sections were incubated with this for 10-20 min. Tissue was then rinsed in 60% ethanol, then four changes of H20. Tissue samples stained with and without oil red O were then stained with hematoxylin and eosin (3 min). The tissue was washed in H20 and mounted in glycerin jelly. Imaging of liver sections was performed on a Nikon Eclipse 50i Brightfield Microscope.

### Immunohistochemistry

Fresh frozen liver tissues preserved in OCT were sectioned (10 μm) and fixed with ice-cold acetone for 5 min. After washing with wash buffer (0.1% tween, 1% BSA in PBS) three times for 5 min, sections were permeabilized (0.25% Triton X-100 and 1% BSA in PBS) for 10 min at room temperature. Tissues were then blocked (0.1% tween, 5% goat serum, 0.5% mouse serum in PBS) for 1 h and stained overnight (4°C) in a humid chamber for α1B-adrenergic receptor (Abcam, ab169523). The following day sections were washed in wash buffer 3 times for 5 min and co-stained for anti-Rabbit IgG Fab2 Alexa Fluor 488 (Cell Signaling, 4412S) at 1:1000 dilution at room temperature for 1 h. Sections were washed 3 times in wash buffer then stained for Alexa Fluor 555 mouse anti-E-cadherin (BD, Cat. 560064) for 1 h at room temperature. Nuclei were stained with DAPI (1:1000 in PBS for 5 min). Imaging of liver sections was performed on BZ-X All-in-One Fluorescence Microscope (Keyence).

### Immunoblotting

Tissue lysates were analyzed by SDS-PAGE, transferred to nitrocellulose membranes, and antibody binding was visualized by enhanced chemiluminescence using a secondary antibody conjugated to horseradish peroxidase (Amersham Biosciences) according to standard procedures. Primary antibodies used were IP_3_R Type 1 (IP_3_R1) (Millipore; Ab5885), IP_3_R Type 2 (IP_3_R2) (Millipore; AB3000), α1B-adrenergic receptor (Abcam, ab169523). The polyclonal anti-SERCA antibodies were generously provided by Dr J. Lytton (University of Calgary, Canada).

### Real time PCR

Total RNA was isolated from hepatocytes using TRIzol reagent followed by column purification (Qiagen), according to the manufacturer’s protocol. DNA contamination was removed with DNAse I treatment (Ampgrade, 1U/µg of RNA, Thermo Fisher Scientific) for 15 min at room temperature. RNA (2 μg) for each sample were simultaneously reverse transcribed using Superscript™ III First-Strand Synthesis System (Thermo Fisher Scientific) according to the manufacturer’s protocol. Quantitative transcript analyses were performed in a AriaMx Real-Time PCR system (Agilent). Optimal conditions were obtained using a five-point, two-fold cDNA and primer dilution curve for each amplicon. Each qPCR reaction contained 12.5 ng of reverse transcribed RNA, each specific primer at 200 nM and SYBR Green PCR Master Mix (Thermo Fisher Scientific), following the manufacturer’s conditions. Relative transcript amount quantification was calculated from three technical replicates, as previously described (Vandesompele *et al*., 2002). Gene expression was normalized to *B2m* (β-2 microglobulin) expression, which did not change under the experimental conditions. Primers are as follows;

Fasn (fatty acid synthase) F 5’-GGAGGTGGTGATAGCCGGTAT-3’, R 5’-TGGGTAATCCATAGAGCCCAG-3’;

G6PC (glucose-6-phosphatase catalytic submit 1) F 5’-TGGCTGGAGTCTTGTCAGG-3’, R 5’-GTAGAATCCAAGCGCGAAAC-3’;

Pepck1 (phosphoenolpyruvate carboxykinase 1) F 5’-ATGTGTGGGCGATGACATT-3’ R 5’-TGGCTGGAGTCTTGTCAGG-3’;

Gnaq (Gq α protein) F 5’-AATCATGTATTCCCACCTAGTCG-3’, R 5’-CCACGAACATTTTCAGGATGA-3’;

Gna11 (G11 α protein) F 5’-CACTGGCATCATCGAGTACC-3’, R 5’-GATCCACTTCCTGCGCTCT-3’;

Plcb1 (Phospholipase Cβ1) F 5’-TTCGTCAAGTGGGATGATGACTC-3’ R 5’-CCACACCTGGCATCCTTGAC-3’;

Plcb3 (Phospholipase Cβ3) F 5’-CGCGGGAGTAAGTTCATCAAA TG-3’, R 5’-CAGTGTGTCCACCTCCATGTT-3’;

Plcb4 (Phospholipase Cβ4) F 5’-GGGAGTAAGTTCATCAAATGGGAC-3’, 5’-AGTGTGTCCACCTCCATGTTG-3’;

Adra1b (α1B-adrenergic receptor) F 5’-TGGGCATTGTAGTCGGAATGT-3’, R 5’-GGGCTTTAGGGTGGAGAACA-3’;

B2m (β2-microglobulin) F 5’-CATGGCTCGCTCGGTGAC-3’, R 5’-CAGTTCAGTATGTTCGGCTTCC-3’.

### Reverse transcriptase PCR to assess X-box binding protein 1 (Xbp1) splicing

cDNA from liver tissue was prepared as described above. PCR was performed using Maxima Hot Start PCR Master Mix (2x) (Thermo #k1061). The PCR conditions were 95 °C 5 min, followed by 25 cycles: 95 °C 30 s, 50 °C 30 s, 72 °C 1 min using an Eppendorf Mastercycler X50 with Xbp1 primers F 5’-ACACGCTTGGGAATGGACAC-3’, R 5’-CCATGGGAAGATGTTCTGGG-3’. Products were resolved on SDS-PAGE gel and the density of spliced (145 bp) and unspliced (171 bp) bands analyzed by Image-J.

### IP_3_ mass assay

Freshly isolated hepatocytes were plated for 1 h, washed and incubated in HBSS for 30 min prior to hormone addition for the required time. The reaction was terminated by aspiration of the HBSS and addition of ice-cold trichloroacetic acid (500 μl). Samples were collected by scraping the tissue culture plates. The precipitated material was removed by centrifugation at 12,000g for 10 min at 4°C, and then 75 μl 10 mM EDTA and 600 μl of tri-*n*-octylamine:1,1,2-trichlorofluoroethane (1:1 ratio) were added to the supernatant and vortexed. The water soluble w phase was collected and the IP_3_ content in the extracted samples was determined using a competition binding assay with [^3^H]-IP_3_ (American Radiochemicals) and IP_3_ binding protein from rat cerebellum (Bredt *et al*., 1989). Extracted samples (100 μl) or IP_3_ standards were added to 100 μl of assay buffer, (50 mM Tris-HCl; 1 mM EDTA; 1 mM dithiothreitol; 1 nM [^3^H]-IP_3_, pH 8.4) and 100 μl of binding protein, and incubated for 1 h at 0°C. The binding protein was collected by centrifugation (12,000g; 10 mins) and aspiration of the supernatant. Bound [^3^H]-IP_3_ was determined by liquid scintillation counting, and the sample IP_3_ determined from the standard curve.

### Serum metabolites

Serum insulin was detected using a sandwich ELISA kit (Thermo Fisher, EMINS) according to the manufacturer’s instructions. Serum cholesterol and ALT/AST levels were determined using a Piccolo® Xpress Chemistry Analyzer (Abaxis) and a Piccolo® Lipid Panel Plus Reagent Disc according to manufacturer’s instructions.

### Statistics

Graph plotting and data analysis was performed with GraphPad Prism software. Statistical analysis was performed using two-tailed students t-test or two-tailed one-way analysis of variance (ANOVA) as indicated in the figure legends.

## Results

### High fat diet model

For this study male C57Bl6 mice were fed 45% HFD or 10% LFD (HFD 58V8, LFD 5755: TestDiet^®^) commencing at 4-5 weeks of age for 1 week and up to 14 weeks. The HFD approximates the Western diet with 30-40% fat (Institute of Medicine (U.S.). Panel on Macronutrients. & Institute of Medicine (U.S.). Standing Committee on the Scientific Evaluation of Dietary Reference Intakes., 2005). We have chosen to study diet-induced obesity rather than use genetic leptin-resistant models such as ob/ob mice, because this more closely resembles disease progression in humans, and data interpretation is not complicated by the systemic disruption of leptin signalling. Our feeding paradigm results in pronounced steatosis after 14 weeks of HFD, but there are no overt histological differences after 1 week (see H&E and Oil Red O staining of liver tissue, Fig. 1). Hyperinsulinemia (plasma insulin measured with ELISA) occurs at both 1 and 14 weeks of HFD (Table 1), and a significant increase in body weight is detected after 8 weeks of HFD compared to the LFD control animals. Increased expression of lipogenic and gluconeogenic genes are a known consequence of hepatic steatosis (Wang *et al*., 2012; Lu *et al*., 2021). The lipogenic gene, fatty acid synthase (FASN), and gluconeogenic genes glucose-6-phosphatase catalytic submit 1 (G6PC) and phosphoenolpyruvate carboxykinase (Pepck1) were all increased after 14 weeks HFD, whereas no changes in gene expression were found after 1 week (Table 1). Table 1 also shows that the serum markers of hepatocyte injury, alanine aminotransferase (ALT) and aspartate transaminase (AST), were both elevated at 14 weeks but not at 1 week of HFD. These data confirm that our feeding paradigm leads to fatty liver disease after prolonged feeding and provides a model for us to investigate the early effects of HFD on the PLC-dependent Ca^2+^ signalling pathway.

**Figure 1.**
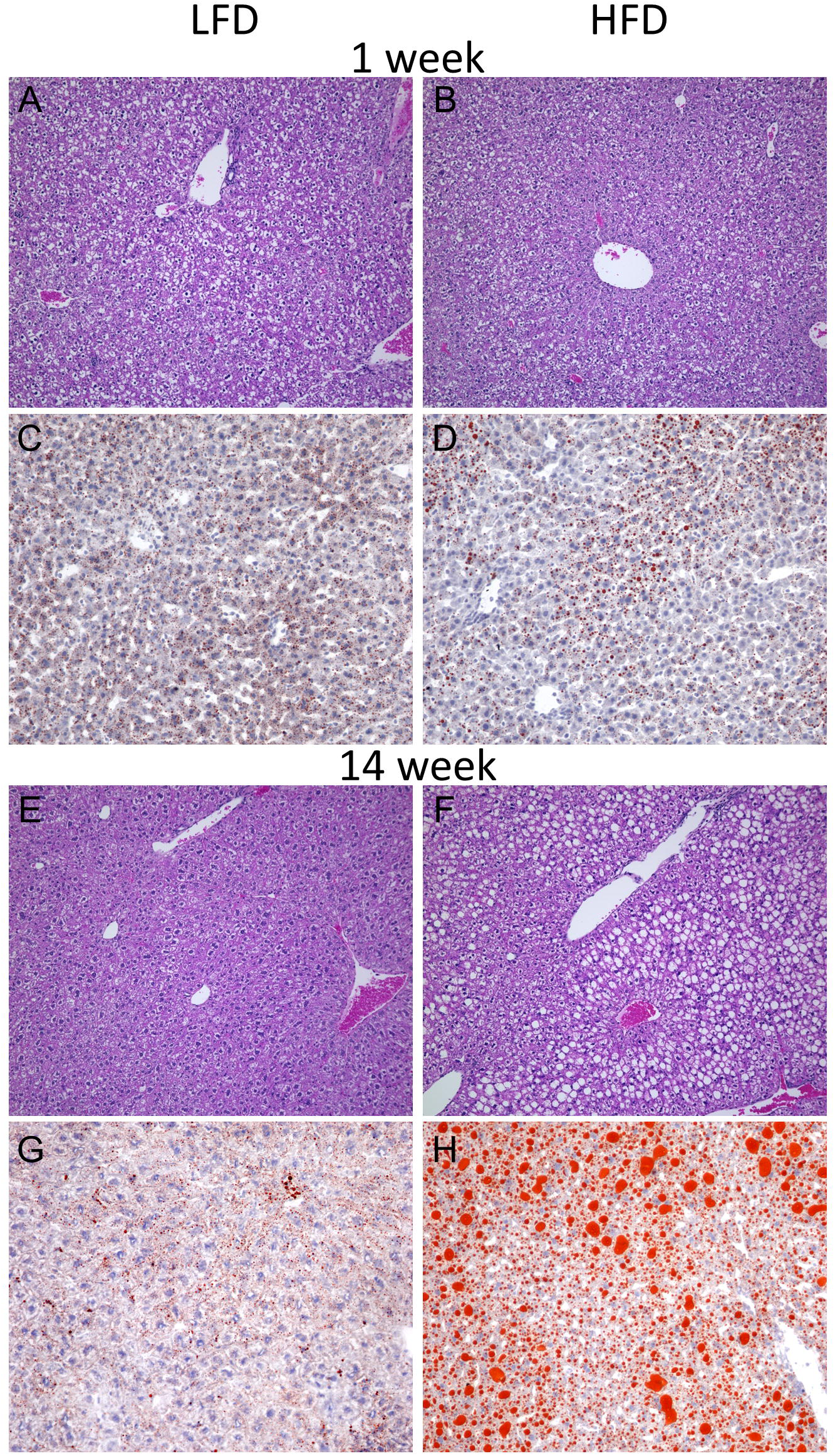
Liver histology of LFD- and HFD-fed mice. Hematoxylin and eosin staining and Oil Red O staining of flash frozen liver sections from mice fed LFD for 1 week (**A, C**) and 14 weeks (**E, G**) or HFD for 1 week (**B, D**) and 14 weeks (**F, H**).

**Table 1.** Blood serum biomarkers and hepatic gene expression changes after HFD feeding. Serum insulin was detected using a sandwich ELISA kit (Thermo Fisher, EMINS) according to the manufacturer’s instructions. Serum cholesterol and ALT/AST levels were determined using a Piccolo® Xpress Chemistry Analyzer (Abaxis). Data are mean ± SD, 4 animals per group. Gene expression changes in fatty acid synthase (Fasn), glucose-6-phosphatase catalytic submit 1 (G6PC) and phosphoenolpyruvate carboxykinase (Pepck1) were determined by RT-PCR normalized to beta-2 microglobulin expression. Data are mean ± SD from ≥ 5 samples per group (*, p<0.05, ***p<0.001 by Student’s unpaired t-test).

|  | **LFD**<br>**(1 week)** | **HFD**<br>**(1 week)** | **p value** | **LFD**<br>**(14 weeks)** | **HFD**<br>**(14 weeks)** | **p value** |
| --- | --- | --- | --- | --- | --- | --- |
| **Serum insulin**<br>**(μIU/ml)** | 15.5 ± 1.7 | 39.2 ± 16.0 | 0.024* | 32.0 ± 10.0 | 54.2 ± 12.1 | 0.039* |
| **Cholesterol**<br>**(mg/dL)** | 99.0 ± 12.9 | 96.8 ± 7.8 | 0.760 | 104.0 ± 9.4 | 188 ± 12.1 | <0.0001*** |
| **ALT U/L** | 35.0 ± 20.8 | 30.6 ± 10.0 | 0.122 | 24.0 ± 5.1 | 62.2 ± 15.6 | 0.003 |
| **AST U/L** | 52.7 ± 16.3 | 60.2 ± 21.7 | 0.589 | 38.7 ± 8.7 | 75.0 ± 13.6 | 0.004 |
| **Fasn** | 1.0 ± 0.8 | 0.91 ± 0.7 | 0.876 | 1.00 ± 0.7 | 1.9 ± 0.4 | 0.042 |
| **G6pc** | 1.0 ± 0.4 | 0.6 ± 0.5 | 0.602 | 1.00 ± 0.5 | 3.6 ± 1.6 | 0.017 |
| **Pepck1** | 1.0 ± 0.4 | 1.0 ± 0.8 | 0.815 | 1.0 ± 0.5 | 1.9 ± 0.6 | 0.044 |

### Short-term HFD attenuates Ca^2+^ signalling

The changes in [Ca^2+^]_c_ elicited by acute stimulation with norepinephrine (NE) were determined in freshly isolated hepatocytes from mice fed LFD or HFD for 1 week (Fig. 2). Hepatocytes loaded with the Ca^2+^ indicator fura-2 were stimulated with a series of increasing doses of NE, followed by a high dose of the purinergic agonist ATP to elicit a maximal [Ca^2+^]_c_ increase. Representative traces of NE-induced [Ca^2+^]_c_ oscillations in individual hepatocytes isolated from LFD and HFD fed animals are shown in Fig. 2A-B. Figs. 2C and 2D show the individual responses of 75 cells from a single coverslip of hepatocytes isolated from LFD and HFD fed animals, respectively, to demonstrate the dose-dependent increases in [Ca^2+^]_c_ responsiveness with sequential additions of increasing NE doses. The traces are presented in a 3-dimension lay out with a colour gradient from blue to red to show the magnitude and frequency of the [Ca^2+^]_c_ response from each hepatocyte (Fig. 2C-D; see scale bar in center). In both LFD and HFD the frequency of [Ca^2+^]_c_ spiking increased with increasing NE dose, and eventually became sustained with the maximal ATP dose. However, the dose response to NE was markedly shifted to the right in the HFD hepatocytes. In the lower dose range from 0.1 to 100 nM NE, more of the LFD hepatocytes responded than HFD hepatocytes, and indeed most hepatocytes from HFD animals did not respond at 10 nM NE and below. At higher doses of NE (100 nM and above), the frequency [Ca^2+^]_c_ oscillations in LFD hepatocytes was markedly higher than that observed for HFD hepatocytes, and the highest does of NE (1 µM) elicited a sustained [Ca^2+^]_c_ increase in many LFD hepatocytes. These changes in [Ca^2+^]_c_ oscillation pattern are investigated further below, but the shift in NE sensitivity across multiple experiments is demonstrated directly by measuring the proportion of hepatocytes responding to each dose of NE (Fig. 2E). This analysis revealed a 10-fold shift in sensitivity between HFD and LFD hepatocytes (EC_50_ ± SEM: LFD 3.06 ± 0.15 nM; HFD 31.33 ± 0.08 nM; n = 6, p < 0.0001).

**Figure 2.**
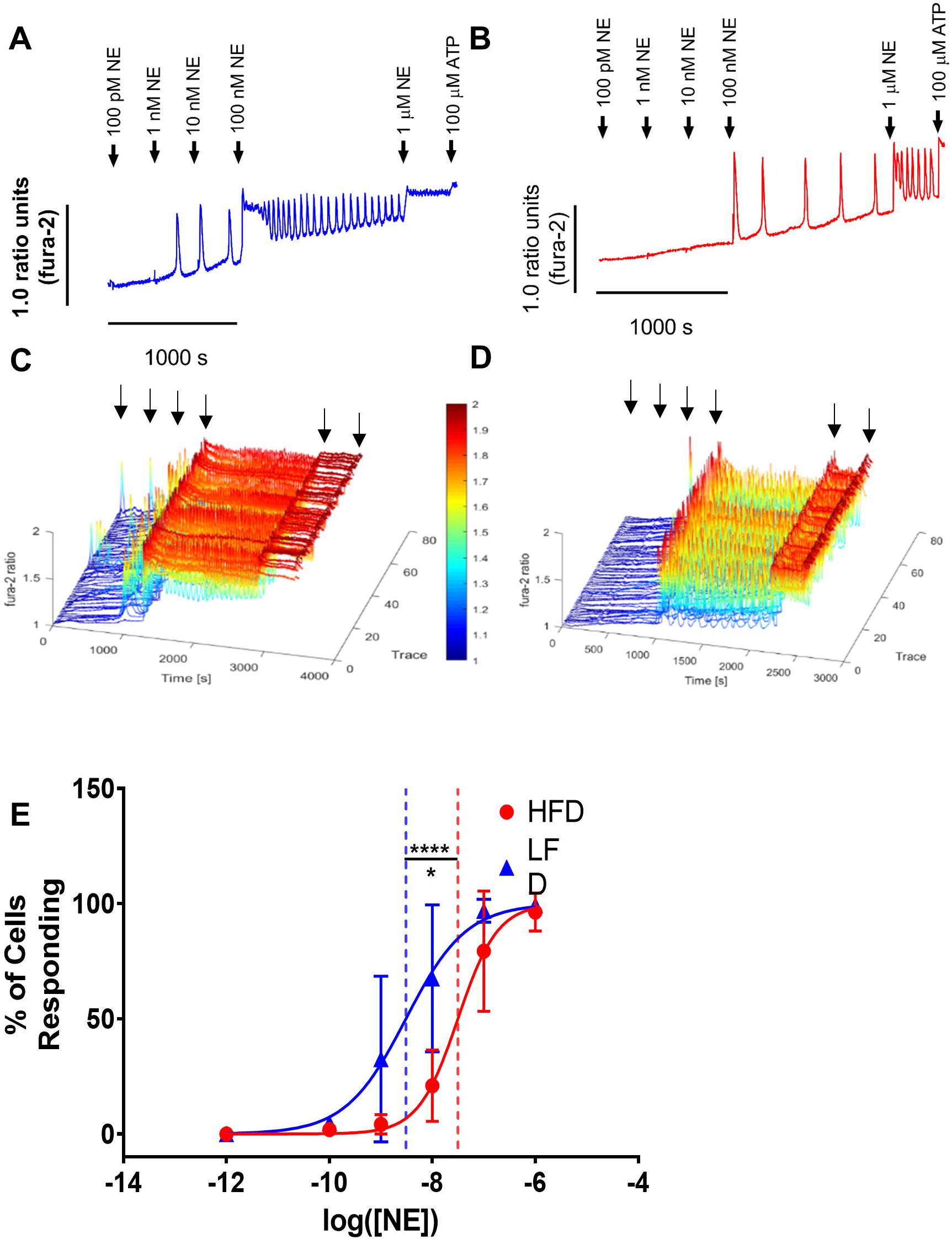
[Ca^2+^]_c_ responses to norepinephrine in mouse hepatocytes. Freshly isolated hepatocytes from mice fed 1 week of HFD or LFD were plated onto collagen-coated glass coverslips and loaded with fura-2 before image time series were recorded with an epifluorescence microscope. Hepatocytes were stimulated with increasing doses of NE as indicated, followed by a maximal dose of ATP. Representative traces of [Ca^2+^]_c_ responses from individual hepatocytes isolated from LFD (**A**) and HFD (**B**) fed mice. (**C-D**) Normalized and colour-mapped view of 75 traces from the same LFD (**C**) and HFD (**D**) animals. (**E**) Proportion of hepatocytes responding to each dose of NE in LFD and HFD animals. Dashed lines represent EC_50_ as calculated by four-parameter normalized dose response fit; log(EC_50_) was significantly different (p <0.0001). Data are plotted as mean ± SD, n = 6 animals per group, >50 cells per animal.

Moreover, significantly more LFD hepatocytes responded to 10 nM NE than HFD hepatocytes (Mean percentage responding ± SD: LFD 67.57 ± 31; HFD 20.91 ± 15.5; n=6, p = 0.009), which reflects the upper range for physiological levels of serum NE (Sharara-Chami *et al*., 2010).

Hepatic metabolism is regulated by the frequency of [Ca^2+^]_c_ oscillations, such that the pattern and duration of [Ca^2+^]_c_ increases are the key determinants of hepatic responsiveness to Ca^2+^-dependent hormones. Therefore, we further quantified the shift in response between hepatocytes isolated from LFD and HFD fed mice by characterizing the type of [Ca^2+^]_c_ response to 100 nM NE, a dose at which a large proportion hepatocytes responded with an increase in [Ca^2+^]_c_ in both groups. [Ca^2+^]_c_ responses from individual hepatocytes were characterized into five groups: no response, single spike, discontinuous oscillations, continuous oscillations and sustained (see Fig. 3A for representative traces of each [Ca^2+^]_c_ signature). The percentage of hepatocytes responding with each [Ca^2+^]_c_ signature from all experiments is shown in Fig. 3B. It is clear that hepatocytes from LFD-fed animals tended towards an oscillatory or sustained response to 100 nM NE, whereas those from HFD-fed animals tended towards less robust responses, including single [Ca^2+^]_c_ spikes and discontinuous [Ca^2+^]_c_ oscillations. This is also visible in the multiple cells projection of Fig. 2C vs 2D. Further analysis of the frequency and width of [Ca^2+^]_c_ spikes in hepatocytes in which oscillations were detected at 100 nM NE revealed that HFD hepatocytes had a significantly lower frequency than LFD cells, with no change in spike width measured as full width half maximum (FWHM) (Fig. 3C and 3D).

**Figure 3.**
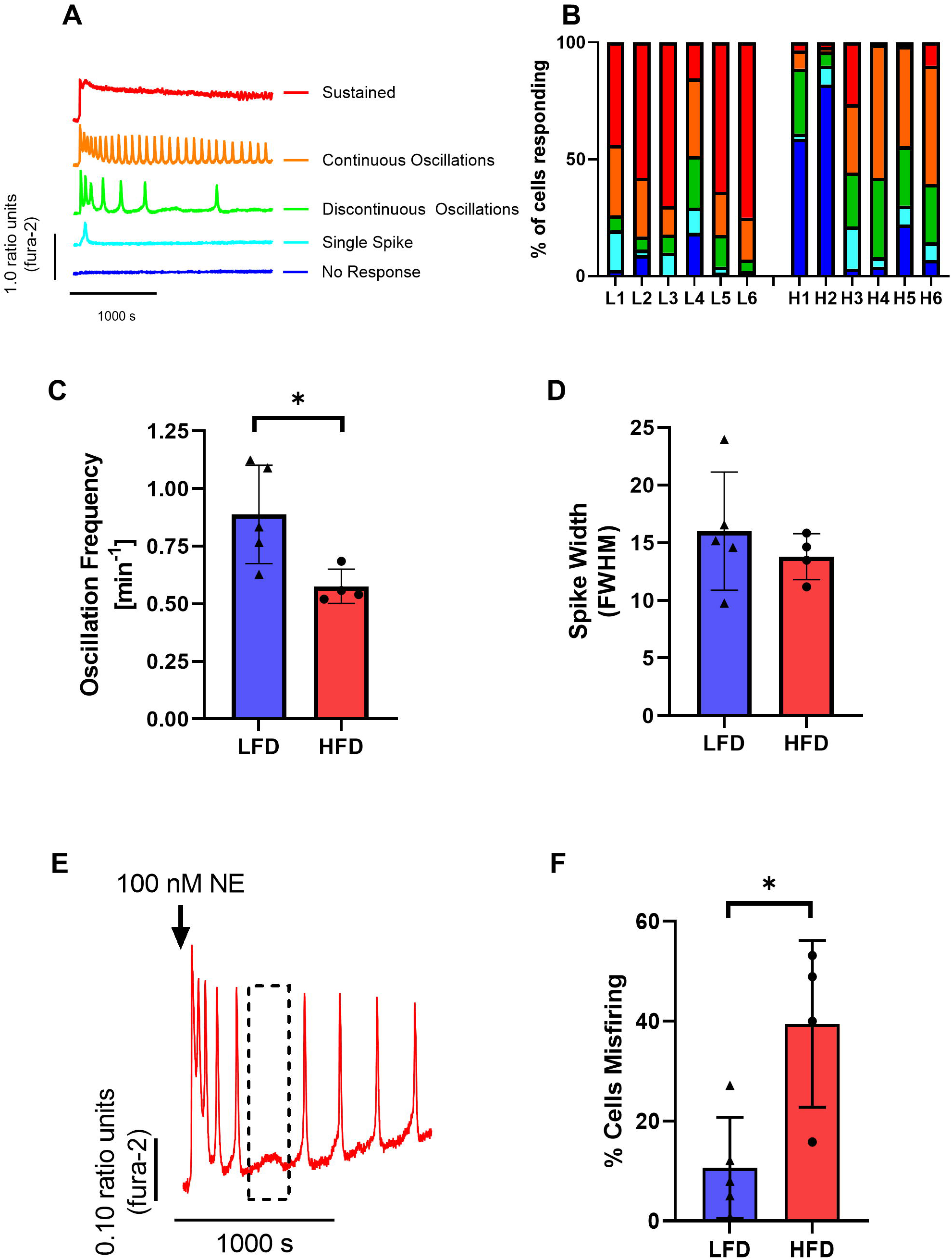
Characterization of [Ca^2+^]_c_ responses to 100 nM norepinephrine. (**A**) Representative [Ca^2+^]_c_ traces for each classification in the ordinal scale used to categorize responses. (**B**) Response type of 6 animals per group (L=LFD, H=HFD). (**C**) Frequency of baseline separate oscillations (hepatocytes with continuous oscillations) to 100 nM NE is reduced after HFD (p = 0.023), but peak width (**D**) is unchanged (p = 0.445). (**E**) Example of a cell misfiring while oscillating. (**F**) A significantly higher proportion of HFD hepatocytes display misfires compared to LFD hepatocytes. Data are plotted as mean ± SD, n = ≥ 4 animals per group. (*, p<0.05, Student’s unpaired t-test).

Not only were hepatocytes from HFD animals less sensitive than LFD hepatocytes to stimulation by NE, the mechanism underlying repetitive [Ca^2+^]_c_ oscillations (Gaspers *et al*., 2014; Bartlett *et al*., 2020) also seemed less robust. During the characterization of the [Ca^2+^]_c_ response patterns, we observed a phenomenon we term “Ca^2+^ spike misfire”, primarily in the HFD cohort. This term describes a failure of a cell to initiate a single [Ca^2+^]_c_ transient in sequence during a train of oscillations, hence Ca^2+^ spike misfire (see Fig. 3E for representative trace). The [Ca^2+^]_c_ spikes elicited by G_αq_/11-coupled receptors in hepatocytes have a biphasic rising behavior (Thomas *et al*., 1991; Gaspers *et al*., 2014), with a slow “pacemaker” phase followed by a more rapid rising phase, due to positive Ca^2+^ feedback on the IP_3_ and on PLC activity (Thomas *et al*., 1991; Gaspers *et al*., 2014; Bartlett *et al*., 2020; Cloete *et al*., 2020). As shown in Fig. 3E, [Ca^2+^]_c_ spike misfires have a pacemaker phase that does not culminate in the generation of a full amplitude [Ca^2+^]_c_ spike, and these typically occur during an ongoing train of [Ca^2+^]_c_ oscillations. Quantification of the number of hepatocytes with [Ca^2+^]_c_ spike misfires revealed that hepatocytes from HFD-fed animals exhibited significantly more misfires than those from LFD-fed animals (Fig. 3F). Taken together, these data demonstrate that short-term (1-week) HFD feeding significantly suppresses NE-induced [Ca^2+^]_c_ signalling, resulting in a decrease in the number of cells that respond, as well as a decrease in the strength of the response elicited.

### Attenuated Ca^2+^ signalling in the intact perfused liver

To further investigate the effect of short-term HFD on the ability of hepatocytes to respond to NE, we investigated Ca^2+^ signalling by intravital imaging in an intact perfused liver preparation (Robb-Gaspers & Thomas, 1995; Gaspers & Thomas, 2005; Bartlett *et al*., 2017; Gaspers *et al*., 2019). To facilitate the intravital measurement of [Ca^2+^]_c_, we generated a hepatocyte-specific GCaMP6f mouse model using the cre-lox system (See Methods). GCaMPf6^hep/cre^ mice were fed 1 week of HFD or LFD prior to surgery and *ex-vivo* perfusion of the liver. Each liver was perfused with increasing concentrations of NE and the changes in hepatocyte [Ca^2+^]_c_ were monitored in real time, with resolution to the single cell level, and fields of view selected to cover 1-3 hepatic lobules. We have shown previously (using livers loaded with fura-2 or fluo3) (Robb-Gaspers & Thomas, 1995; Thomas *et al*., 1995; Gaspers & Thomas, 2005; Bartlett *et al*., 2017; Gaspers *et al*., 2019) that perfusion of the liver with Ca^2+^-dependent hormones elicits oscillatory waves of [Ca^2+^]_c_ increase that propagate through the hepatocytes of entire hepatic lobules, initiating in the periportal (PP) zone and propagating to the pericentral (PC) zone. Confocal images (20x magnification) of the start, peak and end of the initial [Ca^2+^]_c_ oscillation to 30 nM NE challenge are shown in Fig. 4A for a liver from a LFD mouse and Fig. 4B for a liver from a HFD mouse (see also movies in Supplemental files). The red overlay superimposed on the greyscale fluorescence images represents the rate of [Ca^2+^]_c_ increase at each pixel and time point calculated by sequential subtraction to yield a differential image series that tracks the location of [Ca^2+^]_c_ changes. The rightmost panels show maximum projection images for the differential image series, thus mapping all cells/regions that responded during 380 s of perfusion with 30 nM NE. The PP and PC zones of the liver were identified by perfusion with fluorescein-conjugated BSA at the end of the experiment, as described previously (Robb-Gaspers & Thomas, 1995; Thomas *et al*., 1995; Gaspers & Thomas, 2005; Bartlett *et al*., 2017).

**Figure 4.**
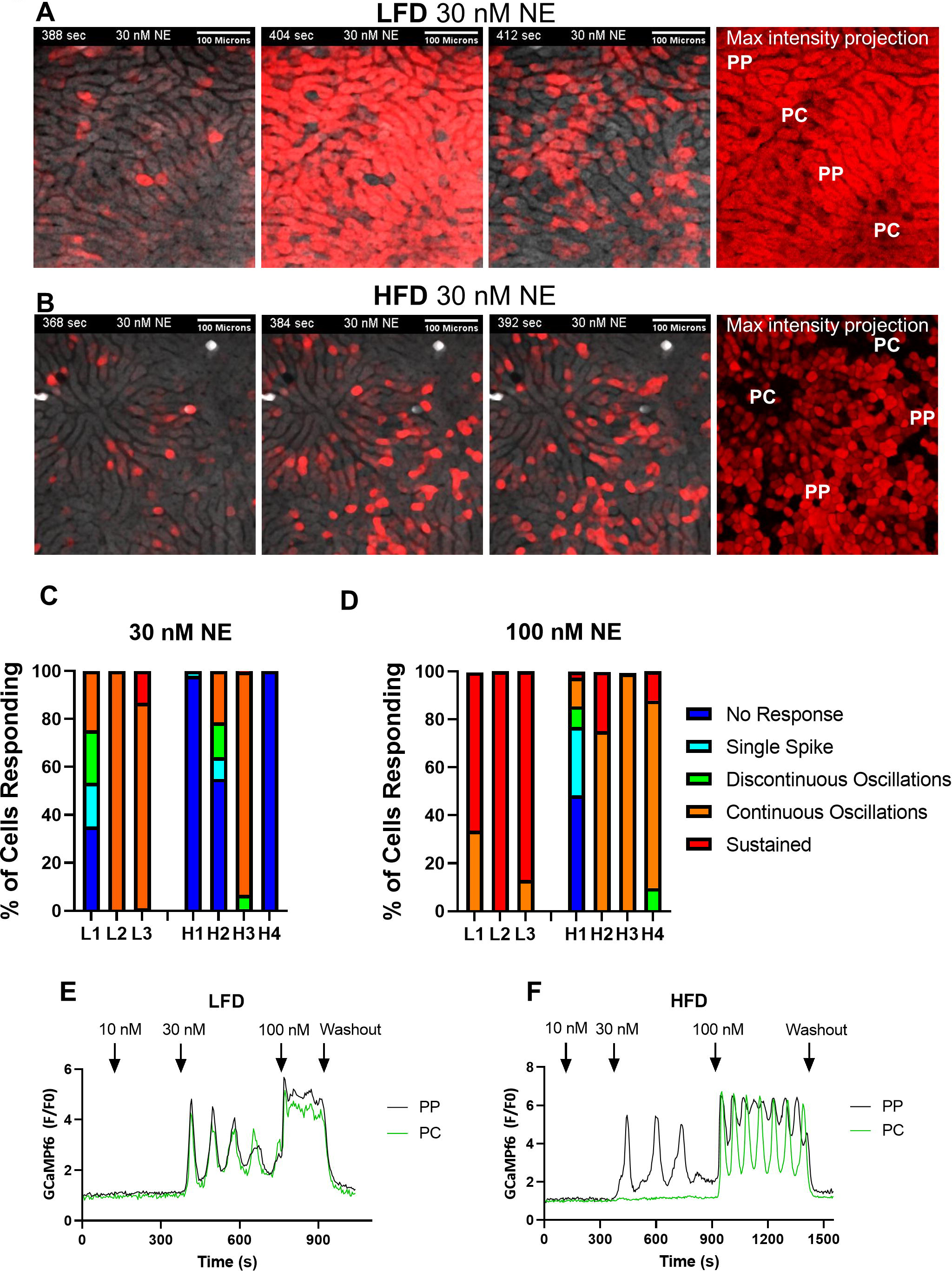
Effect of HFD on norepinephrine-stimulated Ca^2+^ signalling in intact liver. Livers from GCaMP6f^hep/cre^ mice fed LFD or HFD for 1 week were perfused with the indicated concentrations of NE. GCaMP6f fluoresce was recorded with a confocal microscope focused 50-100 µm into the tissue to acquire a [Ca^2+^]_c_ image time series, shown in greyscale. The red overlay images represent the fluorescence change at each time point, calculated by sequential image subtraction to create a differential image series showing the location of [Ca^2+^]_c_ increases for each image frame. **A**) Selected images from a time series at the indicated time points from a LFD liver perfused with 30 nM NE starting 372 s into the perfusion period. Images show the start of the [Ca^2+^]_c_ response (388 s), the peak (404 s) and end (412 s) of the first Ca^2+^ wave for the LFD liver. **B**) Images from a HFD liver perfused with 30 nM NE starting at 352 s, with time points selected to match those in panel A (368 s, 384 s and 392 s) (See also supplementary movies). Maximum intensity projection images in the right panels show the highest value at each pixel from the differential image series, which maps the responsive hepatocytes during a time-matched period (380 s) of 30 nM NE stimulation. Periportal (PP) and pericentral (PC) regions were defined by perfusion with BSA-conjugated fluorescein at the end of each experiment. **C-D**) Quantitation of the type of Ca^2+^ response elicited by 30 nM NE (**C**) and 100 nM NE (**D**) in LFD and HFD GCaMP6f^hep/cre^ livers. Data are mean ± SD from ≥ 3 mice per group. **E-F**) Representative traces of [Ca^2+^]_c_ responses from individual hepatocytes in the PP and PC zones from LFD (**E**) and HFD (**F**) livers sequentially exposed to 10 nM, 30 nM and 100 nM NE.

The number of hepatocytes responding and categorization of individual cell [Ca^2+^]_c_ response patterns to 30 nM NE (Fig. 4C) and 100 nM NE (Fig. 4D) is quantitated in livers from at least 3 animals in each group using the [Ca^2+^]_c_ signatures defined in Fig. 3. The livers from animals fed HFD were less responsive overall than the LFD-fed controls, consistent with the isolated hepatocyte [Ca^2+^]_c_ data (Figs. 2 and 3). Moreover, we observed an impairment in the ability of intercellular [Ca^2+^]_c_ waves to propagate across the liver lobules. This is clearly apparent in the maximum intensity projection images in the rightmost panels of Figs. 4A and 4B, which map the total responsiveness to 30 nM NE in LFD- and HFD-fed livers across multiple lobules. In the LFD livers, [Ca^2+^]_c_ increases occur throughout the entire lobule, reflecting the robust propagation of the NE-induced Ca^2+^ signals. However, the absence of red intensity in the PC regions of the HFD liver indicate that no [Ca^2+^]_c_ increase was observed in these regions throughout the entire perfusion period with 30 nM NE. Nevertheless, these lobular zones were not completely insensitive to NE, because most of the unresponsive cells at 30 nM NE did respond to 100 nM NE, and at this higher NE dose intercellular [Ca^2+^]_c_ waves between hepatocytes were observed to propagate into the PC zone in the HFD livers (see Supplemental movie files). Representative [Ca^2+^]_c_ traces of PP (black) and PC (green) hepatocytes from an individual lobule are shown for a LFD liver (Fig. 4E) and HFD liver (Fig. 4F). In the LFD liver, each [Ca^2+^]_c_ oscillation successfully propagates from the PP to the PC hepatocyte, and they respond with the same coordinated [Ca^2+^]_c_ signature; oscillations at 30 nM NE and sustained at 100 nM NE. By contrast, in the HFD liver lobule, [Ca^2+^]_c_ oscillations are observed at 30 nM NE in the PP but not PC hepatocyte, and [Ca^2+^]_c_ oscillations only propagated through the lobule at 100 nM NE. These data are consistent with the single cell imaging experiments (Figs. 2-3) and demonstrate that HFD impairs NE-dependent Ca^2+^ signal transduction in hepatocytes of the intact liver.

### The expression and distribution of the α1B-adrenergic receptor is unaffected by HFD

One potential mechanism for the reduction in NE-induced Ca^2+^ signalling and intralobular [Ca^2+^]_c_ wave propagation after HFD feeding could be decreased α_1B_-adrenergic receptor expression, or changes in the lobular distribution of the receptor. The distribution of α_1B_-adrenergic receptors in liver was assessed in tissue from LFD (Fig. 5A-C) and HFD (Fig. 5 D-F) animals, and total expression levels were determined by western blotting in tissue lysates samples (Fig. 5G). The location of the α_1B_-adrenergic receptor was not affected by diet (Fig. 5A,D), The PP regions were identified with an established PP protein marker, E-Cadherin (Ben-Moshe *et al*., 2019) (Fig. 5B,E). An overlay of α_1B_-adrenergic receptor (green), E-Cahderin (red) and dapi nuclear stain (blue) is shown in Fig. 5C and 5F. We also detected no change in receptor density in liver tissue lysates (Fig 5G; LFD 47 ± 6 and HFD 53.7 ± 2.6, p = 0.623, data are mean ± SD of protein density normalized to Ponceau stain). These data suggest that the effect of HFD on [Ca^2+^]_c_ signal generation is downstream of the α-adrenergic receptor.

**Figure 5.**
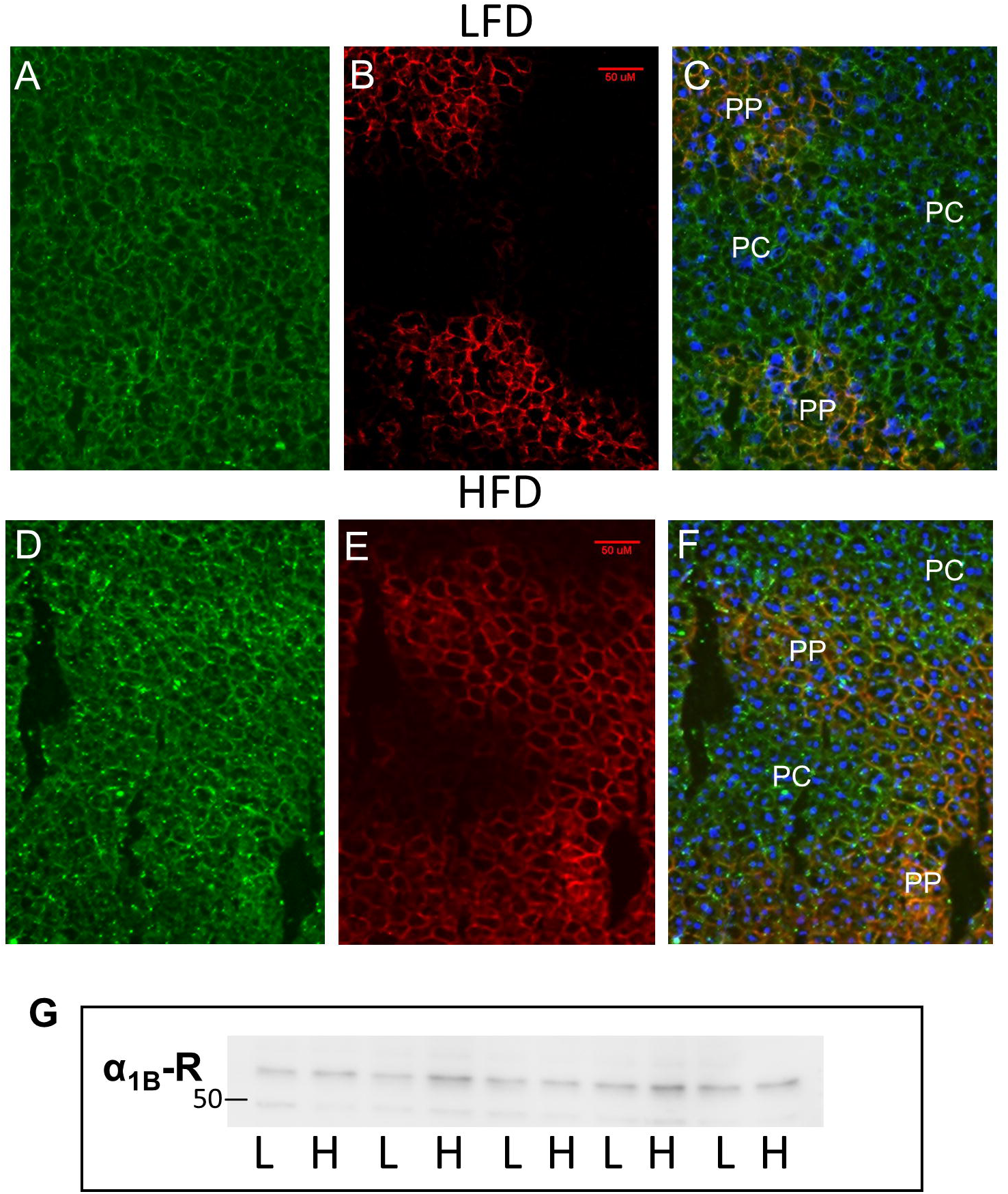
HFD feeding does not alter α_1B_-adrenergic receptor density or lobular distribution. Tissue sections from LFD (**A,B,C**) and HFD (**D,E,F**) mice show the localization of the α_1B_-adrenergic receptor (**A,D**), the periportal marker E-cadherin, scale bar 50 µm.(**D, E**) and an overlay including the nuclear stain dapi (**C,F**). **G**) Western blot of α_1B_-adrenergic receptor from liver tissue lysates of mice fed LFD and HFD for 1 week.

### Short-term HFD does not alter ER Ca^2+^ homeostasis

Previous studies have shown that SERCA activity is decreased in livers from the ob/ob genetic mouse model of obesity (Fu *et al*., 2011). SERCA activity maintains the ER Ca^2+^ store and loss of SERCA activity was shown to correlate with increased ER stress and activation of the unfolded protein response (UPR) (Fu *et al*., 2011). A reduction in SOCE has also been indicated as a factor in the dysregulation of glucose metabolism and development of hepatic steatosis (Wilson *et al*., 2015; Arruda *et al*., 2017). Therefore, it was important to examine whether changes in SERCA activity or SOCE contribute to the Ca^2+^ signalling defects we observe. To determine whether HFD affected ER Ca^2+^, we used the SERCA pump inhibitor thapsigargin (in the absence of extracellular Ca^2+^) to block the pump and release the ER Ca^2+^ pool. These studies revealed no change in ER Ca^2+^ load between hepatocytes isolated from LFD or HFD mice (Fig. 6A, representative traces; Fig. 6B, summary data). We also assessed the rate of plasma membrane Ca^2+^ influx via SOCE by adding Ca^2+^ back to the extracellular buffer after complete emptying of the ER Ca^2+^ stores with thapsigargin (Figs. 6A). This protocol also allowed us to measure rates of Ca^2+^ efflux from the cell via the plasma membrane Ca^2+^-ATPase pump by subsequently removing the extracellular Ca^2+^. There was no effect of HFD on either SOCE or on the rate of Ca^2+^ extrusion from the cell (Fig. 6C). Moreover, there was no change in SERCA expression level between the two groups (Fig. 6D). ER stress and the UPR induced by ER Ca^2+^ depletion leads to splicing of X-box binding protein-1 (Xbp1) mRNA (van Schadewijk *et al*., 2012). However, Xbp1 mRNA revealed no shift in the spliced/unspliced distribution of Xbp1 in hepatocytes from HFD mice compared to LFD mice (Fig. 6E). Together these data show that 1 week of HFD feeding does not impair hepatocyte Ca^2+^ homeostasis or induce ER stress.

**Figure 6.**
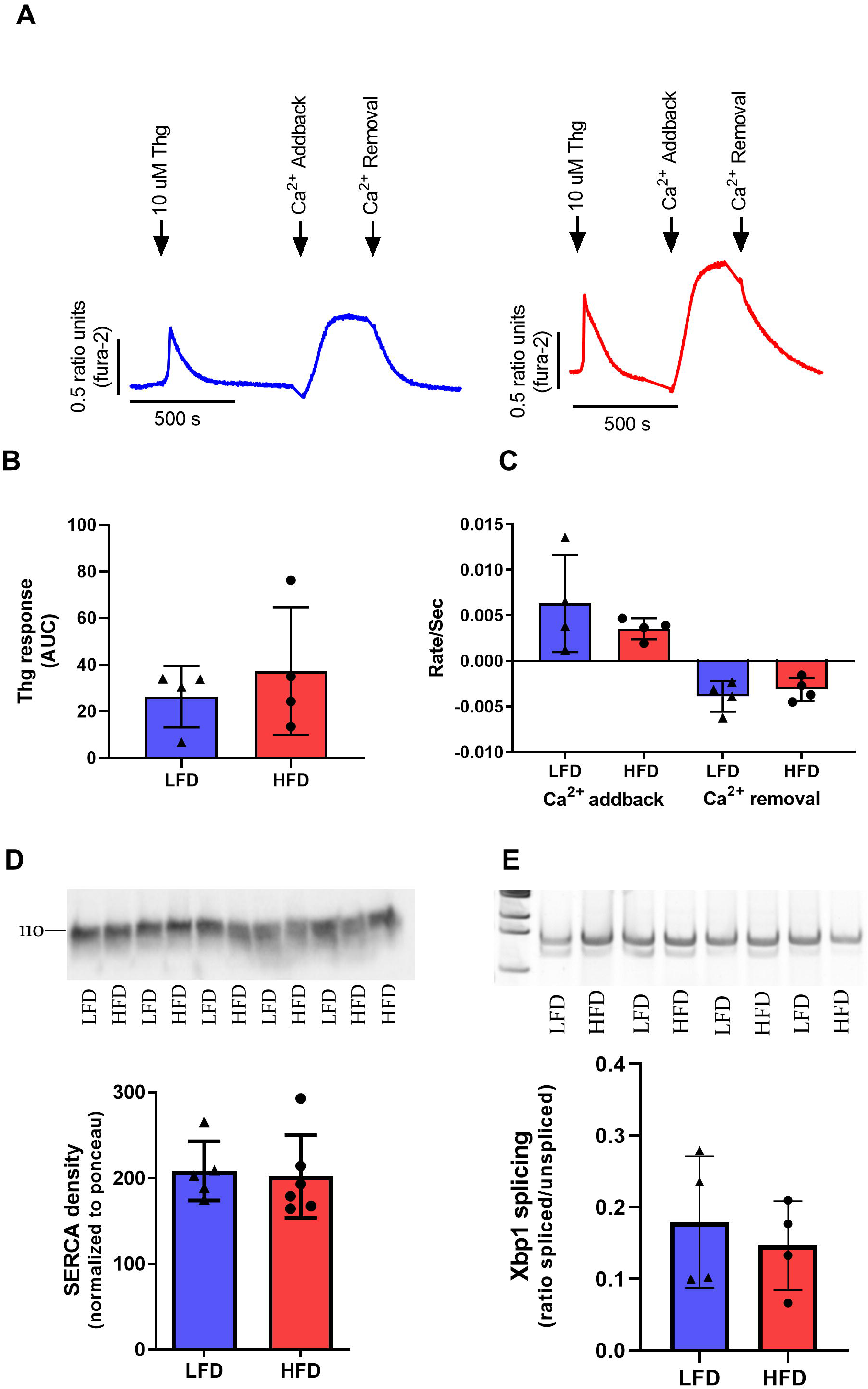
HFD does not impair ER Ca^2+^ homeostasis. Freshly isolated hepatocytes were loaded with fura-2AM then treated with thapsigargin (Thg) in Ca^2+^-free buffer, followed byCa^2+^ addback, and then removal. (**A**) Representative traces of LFD (blue) and HFD (red) fed animals. Area under curve (AUC) of thapsigargin response, p = 0.496(**B**), and rates of Ca^2+^ influx following Ca^2+^ addback (p = 0.348) and efflux (p = 0.492) after subsequent removal of extracellular Ca^2+^ (**C**). Data are mean ± SD from 4 animals per group. **D**) Western blot and densitometry of SERCA in tissue lysates from mice fed LFD and HFD for 1 week (mean ± SD from ≥ 5 animals per group, p =0.805). **E**) Agarose gel and quantification of Xbp1 splicing in isolated hepatocytes from mice fed LFD and HFD for 1 week (mean ± SD from 4 animals per group, p = 0.576).

### No change in IP_3_ receptor function after HFD feeding

We next turned our attention to the IP_3_Rs to determine whether changes in channel activity were involved in the observed Ca^2+^ signalling defect following HFD. Several groups have reported changes in IP_3_R expression and function in other mouse models, although these are not always consistent (Wang *et al*., 2012; Arruda *et al*., 2014; Feriod *et al*., 2014; Feriod *et al*., 2017; Khamphaya *et al*., 2018). Therefore, we performed a series of experiments to assess potential changes in the abundance and activity of the hepatic IP_3_Rs in response to LFD and HFD. Western blot of tissue membrane lysates revealed no significant difference in expression of either IP_3_R1 or IP_3_R2 after 1 week of feeding (Fig. 7A; summary data shown in Fig. 7B). To assess IP_3_R function we employed a permeabilized cell assay to allow the direct activation of IP_3_Rs and measured the dose-response and maximum Ca^2+^ release with exogenously applied IP_3_. Freshly isolated hepatocytes were loaded with the low affinity Ca^2+^ indicator fura-2FF/AM for 1 h to allow dye compartmentalization into the ER. The hepatocyte membrane was then permeabilized with digitonin to release cytosolic dye, which enabled us to measure changes in ER luminal [Ca^2+^] ([Ca^2+^]_ER_) (see Methods) (Hajnóczky *et al*., 1994; Hajnóczky & Thomas, 1997; Gaspers *et al*., 2021). After permeabilization, the cells were incubated with ATP to load the ER with Ca^2+^ (Fig. 7C). This was followed by a test dose of IP_3_, a maximal dose of IP_3_, and finally ionomycin to release all compartmentalized Ca^2+^ and normalize for plating density and dye load. Summary data for the IP_3_ dose-response in hepatocytes isolated from mice fed LFD and HFD for 1 week (n=4 for each group) are shown in Fig. 7D. We found no significant difference between LFD and HFD in the maximum amplitude of [Ca^2+^]_ER_ decrease (Mean ± SD; LFD 0.33 ± 0.04 and HFD 0.37 ± 0.04 change in fura2-FF ratio, p = 0.347, n = 3), or in the EC_50_ for IP_3_ (Mean ± SEM, LFD 234 ± 20 nM and HFD 218 ± 13 nM, p = 0.887, n = 3).

**Figure 7.**
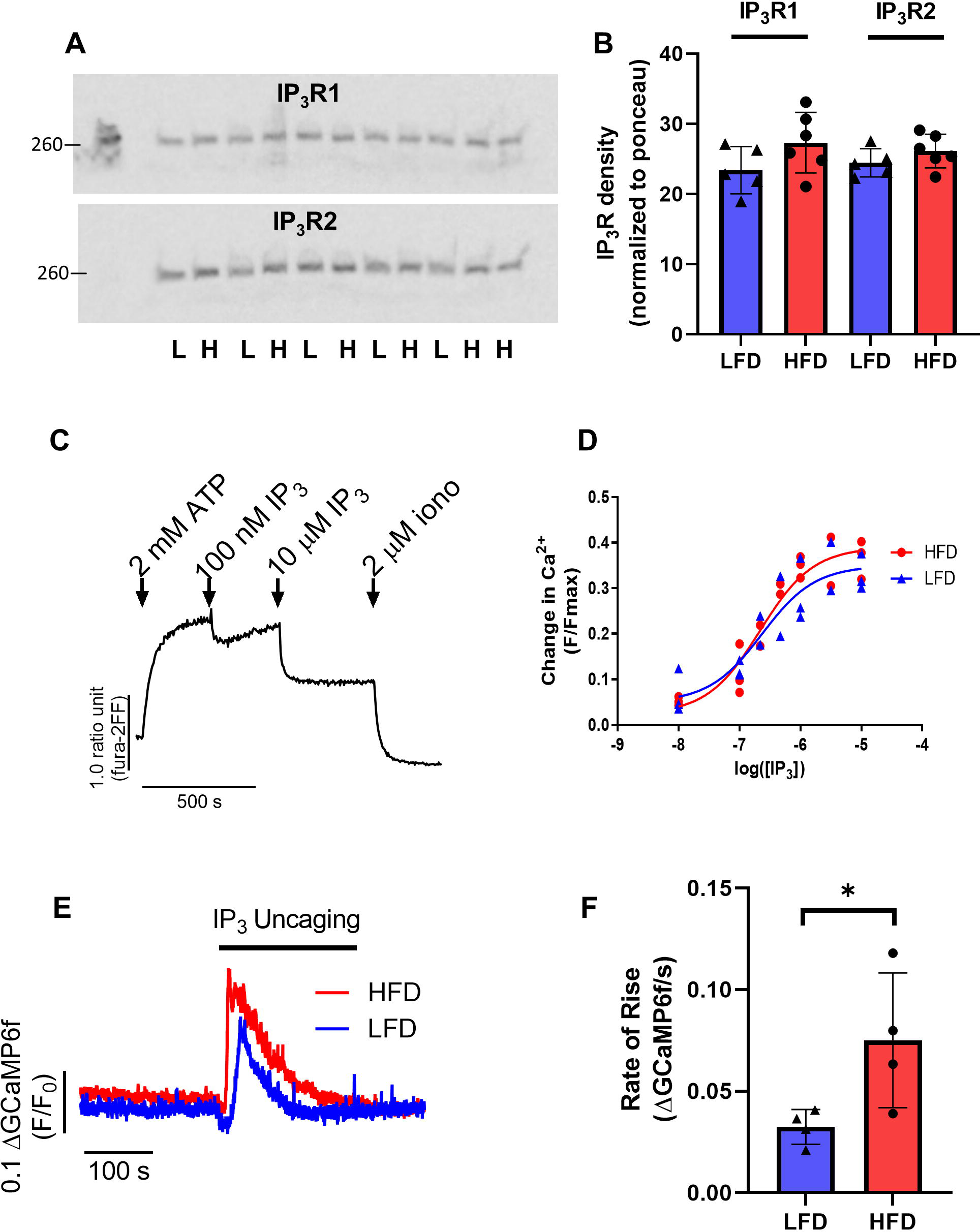
Effect of short-term HFD feeding on IP_3_ receptor function. **A)** Western blot and **B**) protein density of IP_3_R1 and IP_3_R2 in membrane fractions of liver tissue lysates (data are mean ± SD of ≥ 5 mice per group, IP_3_R1 p = 0.135; IP_3_R2 p = 0.247). **C-D**) Freshly isolated hepatocytes were loaded for 1 h with fura-2FF, and then the plasma membrane was permeabilized with 25 µg/ml digitonin for 10 min to allow measurements of [Ca^2+^]_ER_ using the compartmentalized fura-2FF. (**C**) Addition of ATP to the permeabilized hepatocytes led to loading of the ER with Ca^2+^ before the cells were stimulated with IP_3_. Changes in [Ca^2+^]_ER_ after IP_3_ addition were normalized to the difference between the peak ATP and the maximum [Ca^2+^]_ER_ decrease after the addition of 2 µM ionomycin (iono), (calcium change F/Fmax). The [Ca^2+^]_ER_ release to each IP_3_ dose was then plotted to obtain dose response curves for hepatocytes from the LFD and HFD mice (n = 4 per group) (**D**). No change in IP_3_ sensitivity was observed after 1 week of HFD (EC_50_ p = 0.608). **E**) Representative traces showing the [Ca^2+^]_c_ response to IP_3_ uncaging in LFD and HFD hepatocytes measured with GCaMP6f. A significantly faster rate of [Ca^2+^]_c_ rise (**F**) was observed in HFD compared to LFD hepatocytes (p = 0.0474, Student’s unpaired t-test). Data are plotted as mean ± SD of 4 mice and >50 cells per group).

One potential caveat with the permeabilized cell approach is that there could be loss of regulatory cytoplasmic components, including kinases and phosphatases that may modulate the sensitivity of the IP_3_Rs *in vivo*. For example, insulin and glucagon signalling cascades respectively decrease and increase the sensitivity of the IP_3_R through site-specific phosphorylation (Khan *et al*., 2006; Betzenhauser *et al*., 2009; Wang *et al*., 2012), which could be lost when cells are permeabilized. Therefore, we used IP_3_-uncaging in intact hepatocytes to investigate whether IP_3_R activity or sensitivity to IP_3_ is modified following 1 week of HFD feeding. This approach enabled us to test the sensitivity of the IP_3_R to IP_3_ independent of the upstream receptor-PLC signalling pathway that generates IP_3_. For this study, we transfected hepatocytes with GCaMP6f and cultured overnight to allow for expression of the indicator (see Methods). This molecular Ca^2+^ indicator buffers [Ca^2+^]_c_ to a lesser degree compared to chemical indicators such as fura-2, making it better suited for detecting small changes in [Ca^2+^]_c_ and fast kinetics. Isolated hepatocytes from LFD- and HFD-fed mice were loaded with caged-IP_3_, then, following an initial baseline recording, stimulated with UV light exposure to continuously release IP_3_, as described previously (Bartlett *et al*., 2015; Bartlett *et al*., 2017; Bartlett *et al*., 2020). We found that hepatocytes from mice fed HFD for 1 week were more sensitive to IP_3_ uncaging compared to LFD controls. Thus, the increase in [Ca^2+^]_c_ evoked by IP_3_ uncaging reached the maximum faster in HFD hepatocytes (see Fig. 7E for representative traces), and had a faster rate of rise (summarized in Fig. 7F). These data demonstrate a sensitization of the IP_3_R population in intact hepatocytes from the HFD mice, which has not been directly observed previously, despite a number of studies suggesting increases in IP_3_R expression and alterations in IP_3_R phosphorylation (Wang *et al*., 2012; Arruda *et al*., 2014; Feriod *et al*., 2017). Nevertheless, this enhanced IP_3_R activity does not explain the reduced responsiveness to hormone after 1 week of HFD exposure, and may represent an adaptation to the early suppression of Ca^2+^ signalling in response to HFD. Thus, the absence of any change in hepatocyte Ca^2+^ homeostasis or decrease in IP_3_R function, suggests that the lesion in the hormone-stimulated Ca^2+^ signalling pathway lies upstream of the IP_3_ receptor.

### PLC activity is significantly attenuated by HFD

We investigated whether changes in IP_3_ generation are responsible for the observed suppression of NE-induced [Ca^2+^]_c_ responses after HFD. Hepatocytes isolated from HFD or LFD mice were plated for 1 h, and then stimulated with 100 nM NE for 15, 30, 45 and 60 s (a time-frame in which IP_3_ levels peak after hormone stimulation) (Bartlett *et al*., 2017). IP_3_ levels were determined using a radiolabeled competition binding assay. Mean IP_3_ production at each time point is shown in Fig. 8A and the mean peak response in Fig. 8B. Peak IP_3_ production was significantly decreased in hepatocytes from mice fed the HFD compared to LFD. This finding indicates a deficit in the ability of the α_1B_-adrenergic receptor to activate PLC after HFD consumption. Gene expression analysis of the receptor, G protein and PLC isoforms involved in the α_1B_-adrenergic signalling cascade revealed no diet-dependent changes in expression (Fig. 8C), suggesting that the signalling deficit is due to decreased PLC activity rather than a change in the expression of any one signalling component. Thus, it appears that an early event in the response of the hepatocyte to a HFD is to suppress Ca^2+^ signalling elicited by physiological levels of IP_3_-dependent hormones, which may be the precipitating factor in subsequent changes in IP_3_R function, expression and coupling to mitochondrial metabolism (Wang *et al*., 2012; Arruda *et al*., 2014; Feriod *et al*., 2017).

**Figure 8.**
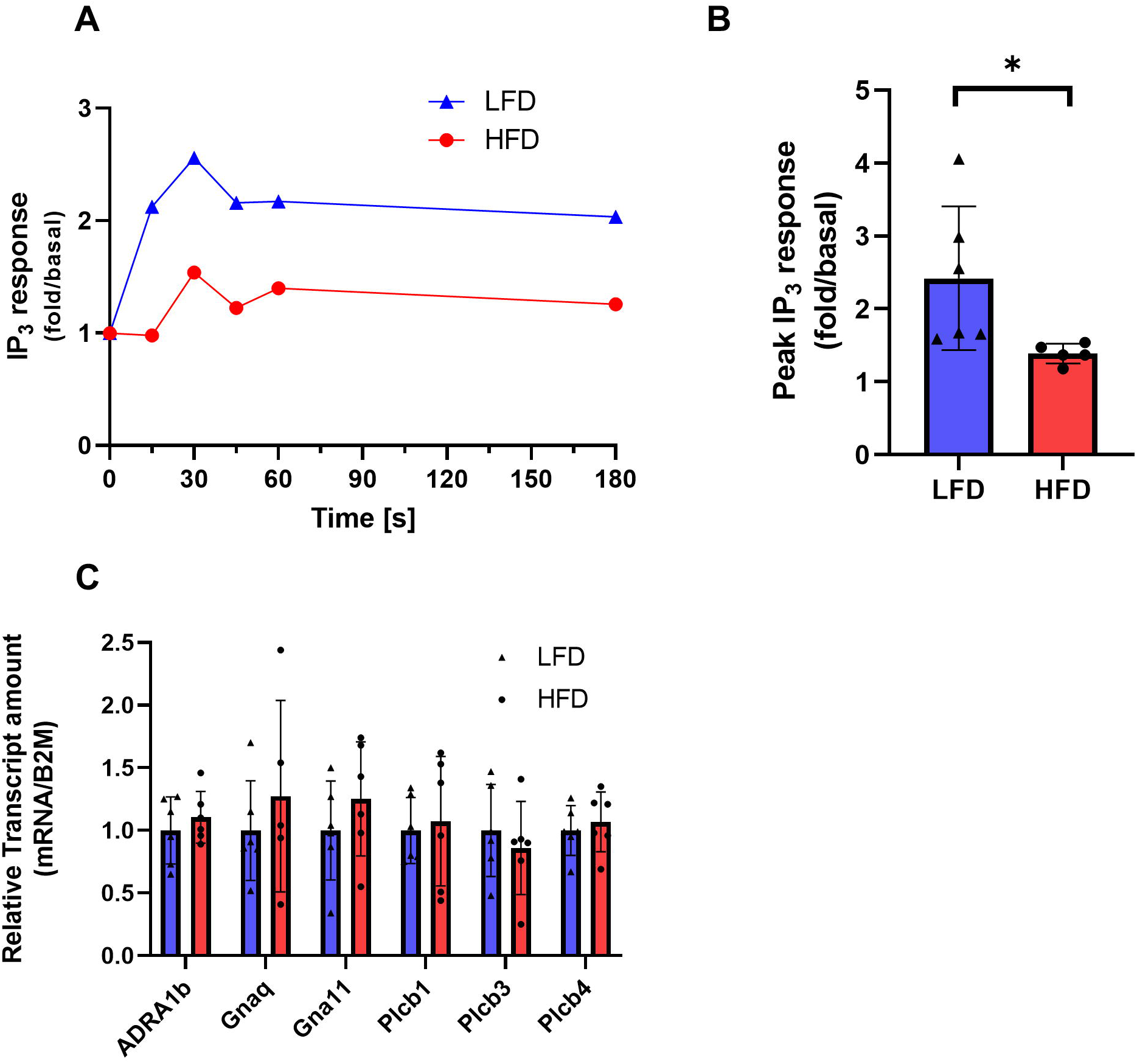
Norepinephrine-induced IP_3_ production is attenuated by short-term HFD. Freshly-isolated hepatocytes were stimulated with 100 nM NE for 15, 30, 45 60 and 180 s. The reaction was terminated with trichloroacetic acid and the water-soluble components extracted for determination of IP_3_ content using a [^3^H]-IP_3_ competition binding assay. **A**) Representative IP_3_ time-course, and **B**) mean peak IP_3_ response to NE stimulation are shown (maximal response time in both groups varied between the 15 and 30 s time points), data are mean ± SD from ≥ 5 animals (* p<0.05, Student’s unpaired t-test). **C**) Comparison of gene expression profiles of the signalling components upstream of IP_3_R, ADRA1b (α_1B_-adrenergic receptor) p = 0.463, Gnaq (G_qα_ protein), p = 0.460, Gna_11_ (G11 α protein) p = 0.330, Plcb (Phospholipase Cβ)1, p = 0.763, 3, p = 0.527 & 4, p = 0.602 (data are mean ± SD, n=≥5).

## Discussion

GPCR-dependent [Ca^2+^]_c_ signalling plays a fundamental role in regulating hepatic metabolism to activate glucogenic signalling pathways and oxidative metabolism, whilst opposing insulin-dependent signalling cascades that lead to glucose storage and lipogenesis (Bartlett *et al*., 2014). Increases in [Ca^2+^]_c_, most often occurring as periodic [Ca^2+^]_c_ oscillations, regulate glucose metabolism by stimulating glycogen phosphorylase activity and negatively regulating glycogen synthase, as well as stimulating *de novo* glucose synthesis by activation of gluconeogenesis (Blackmore *et al*., 1986; Kraus-Friedmann & Feng, 1996; Bartlett *et al*., 2014). These Ca^2+^-dependent hormones also regulate lipid metabolism by inhibiting acetyl-coA-carboxylase, suppressing fatty acid synthesis and promoting β-oxidation; again pathways that antagonize the anabolic effects of insulin (Brownsey *et al*., 2006). Therefore, diet-induced changes in hormonal Ca^2+^ signalling have the potential to significantly alter hepatic metabolism and energy homeostasis. However, despite the essential role of α_1B_-adreneric receptor-mediated signalling in maintaining glycemic control and hepatic carbohydrate and lipid homeostasis, this is the first study to determine the effects of HFD on [Ca^2+^]_c_ signal generation and transduction at the cellular and organ level. Our data show that Ca^2+^ signalling in response to NE is impaired by short-term HFD feeding. This inhibition results from attenuation of NE-induced IP_3_ generation, which in turn leads to suppression of [Ca^2+^]_c_ increases, manifest as reduced Ca^2+^ oscillation frequency and cellular responsiveness, as well as decreased propagation of intralobular Ca^2+^ waves in the intact liver.

Our previous studies have delineated the positive and negative feedback mechanisms that allow for baseline-separated [Ca^2+^]_c_ oscillations, such as those observed in hepatocytes (Gaspers *et al*., 2014; Bartlett *et al*., 2015; Bartlett *et al*., 2020; Cloete *et al*., 2020). Biphasic Ca^2+^ regulation of IP_3_Rs is well established, with elevation of [Ca^2+^]_c_ increasing open probability and eliciting Ca^2+^-induced Ca^2+^ release, which is followed by inhibition of IP_3_R channel activity at high [Ca^2+^]_c_, giving rise to periodic coordinated bursts of Ca^2+^ release and reuptake (Bezprozvanny *et al*., 1991; Foskett *et al*., 2007). We have demonstrated that an addition essential component of baseline-separated [Ca^2+^]_c_ oscillations is positive feedback of Ca^2+^ on PLC to create regenerative cycles of IP_3_ production and Ca^2+^ release (Politi *et al*., 2006; Gaspers *et al*., 2014; Thurley *et al*., 2014; Bartlett *et al*., 2015; Bartlett *et al*., 2020; Cloete *et al*., 2020). This feedback to increase PLC activity ensures each periodic rise in IP_3_ is sufficient to elicit a full amplitude [Ca^2+^]_c_ spike and drive intra- and intercellular propagation of robust spatially distributed [Ca^2+^]_c_ oscillations and waves (Gaspers *et al*., 2014; Bartlett *et al*., 2020; Cloete *et al*., 2020). These mechanisms ensure that even the low [Ca^2+^]_c_ oscillation frequencies induced by submaximal physiological doses of IP_3_-linked hormones can fully penetrate the hepatocyte to regulate hepatic metabolism and mitochondrial function (Hajnoczky *et al*., 1995; Robb-Gaspers *et al*., 1998a; Robb-Gaspers *et al*., 1998b; Jouaville *et al*., 1999; Denton, 2009; Cardenas *et al*., 2010). Protein Kinase C also plays a key role in the generation of baseline-separated [Ca^2+^]_c_ oscillations (Sanchez-Bueno *et al*., 1990; Berrie & Cobbold, 1995; Bartlett *et al*., 2015; Cloete *et al*., 2021; Corrêa-Velloso *et al*., 2021). Of particular relevance to the present work, our previous studies have shown that PKC inhibits IP_3_ formation and reduces the frequency of hormone-induced [Ca^2+^]_c_ oscillations (Bartlett *et al*., 2005; Bartlett *et al*., 2015; Cloete *et al*., 2021; Corrêa-Velloso *et al*., 2021). We propose that HFD modifies these essential feedback loops, giving rise to impaired [Ca^2+^]_c_ signalling in hepatocytes.

Our data show that HFD reduces the number of hepatocytes responding to NE (Fig. 2) and also shifts the Ca^2+^ signatures to less robust responses and lower [Ca^2+^]_c_ oscillation frequencies (Fig. 3). Individual [Ca^2+^]_c_ spikes are comprised of a pacemaker period during which there is a slow rise in [Ca^2+^]_c_, followed by a rapid rising phase due to the positive Ca^2+^ feedback on PLC (Thomas *et al*., 1991; Gaspers *et al*., 2014). In addition to the overall reduction in [Ca^2+^]_c_ oscillation frequency, we also observed a propensity for skipped [Ca^2+^]_c_ oscillations in hepatocytes isolated from HFD-fed mice, whereby the pacemaker phase occurs but fails to elicit the rapid whole cell rise in [Ca^2+^]_c_. Together with the overall suppression of Ca^2+^ signalling, this implies a reduction in PLC activity either due to inhibition of the enzyme itself or reduced coupling between the α_1B_-adreneric receptor and PLC. We observed no changes in α_1B_-adreneric receptor density or lobular distribution, and this led us to determine whether HFD resulted in reduced PLC activity. Specifically, we carried out biochemical measurements of IP_3_, which led to our novel finding that short-term consumption of HFD impairs NE-induced IP_3_ generation in hepatocytes. As discussed below, we suggest that the IP_3_ response to NE is dampened in HFD hepatocytes as a result of an imbalance in the feedback loops that maintain [Ca^2+^]_c_ oscillations.

Intact perfused liver experiments also showed that HFD suppresses Ca^2+^ signalling in hepatocytes *in situ*, and further demonstrated that this is associated with a reduction in the radial spread of intercellular [Ca^2+^]_c_ wave propagation across the liver lobules during perfusion with NE (Fig. 4). As we have described previously (Robb-Gaspers *et al*., 1998a; Bartlett *et al*., 2014; Gaspers *et al*., 2019), the intralobular [Ca^2+^]_c_ waves initiate in a subset of periportal cells that are most sensitive to NE and other Ca^2+^-dependent hormones. Trans-lobular intercellular [Ca^2+^]_c_ waves propagate as a result of Ca^2+^ and IP_3_ transfer through gap junctions of adjacent hepatocytes, but also rely on the systemic presence of the hormone throughout the lobule (Robb-Gaspers & Thomas, 1995; Thomas *et al*., 1995; Gaspers & Thomas, 2005). Thus, under normal conditions, perfusion with hormone or catecholamine results in coordinated [Ca^2+^]_c_ signals being propagated across the entire liver lobule from periportal to pericentral zone hepatocytes. It is possible that the disruption of this integrated signalling in response to HFD could reflect impaired hepatic gap junction intercellular communication. However, full lobular Ca^2+^ wave propagation was observed at higher NE doses in livers from HFD mice, suggesting that gap junction permeability is unlikely to be the limiting factor. Thus, these data are more consistent with the suppression of IP_3_ formation observed in isolated HFD hepatocytes, which would effectively reduce the ability to propagate regenerative Ca^2+^ waves between all of the hepatocytes across the lobule.

As noted above, PKC plays a key role as a negative regulator of GPCR-stimulated IP_3_ formation (Bartlett *et al*., 2015; Corrêa-Velloso *et al*., 2021). Thus, one potential mechanism for the suppression of Ca^2+^ signalling in hepatocytes from mice fed a HFD is a higher level of negative-feedback by PKC. Significantly, elevated PKC activity is associated with diet-induced obesity. Oversupply of free fatty acids and triacylglycerol lead to elevated intracellular DAG, which increases nPKC activity (Geraldes & King, 2010; Greene *et al*., 2010; Schmitz-Peiffer & Biden, 2010; Kumashiro *et al*., 2011; Turban & Hajduch, 2011). One established consequence of this constitutive PKC activation is insulin resistance due to PKC-mediated phosphorylation and inactivation of IRS-1 & 2 (Itani *et al*., 2002; Yu *et al*., 2002; Boden *et al*., 2005; Samuel *et al*., 2007; Ritter *et al*., 2015). GPCRs and PLCβ are also known PKC targets (Alcántara-Hernández *et al*., 2000; Strassheim & Williams, 2000; Young *et al*., 2003). It is therefore possible that persistent nPKC activation leads to a tonic inhibition of agonist-stimulated IP_3_ formation, over and above the periodic negative feedback by DAG that is generated directly by the GPCR/PLC complex itself. Given the strong evidence that PKC enzymes negatively regulate [Ca^2+^]_c_ signals and IP_3_ generation (Bartlett *et al*., 2015; Corrêa-Velloso *et al*., 2021), we propose that this is a primary mechanism by which HFD impairs Ca^2+^ signalling in the liver. Insulin resistance is a well-established response to HFD, and our data suggest that the opposing catabolic signalling pathways activated by NE are also inhibited as a result of HFD. One potential consequence of this would be reduced lipolysis and fatty acid oxidation, which could contribute to the accumulation of intrahepatic triglycerides that are associated with metabolic syndrome, despite the insulin-resistant state.

In the last several years a number of studies have investigated the relationship between obesity and altered hepatic Ca^2+^ homeostasis, predominantly using genetic obesity models. Decreased SERCA activity has been shown to reduce ER Ca^2+^ load, leading to ER stress and unfolded protein response, whereas SERCA overexpression reversed ER stress and improved glucose homeostasis in these animal models (Park *et al*., 2010; Fu *et al*., 2011; Zhang *et al*., 2014). A reduction in SOCE has also been observed in ob/ob mice (Arruda *et al*., 2017) and obese Zucker rats (Wilson *et al*., 2015). Incubation of isolated hepatocytes from lean rats with amiodarone or palmitate to increase intracellular lipid droplets also impaired SOCE, indicating that lipid accumulation mediates this effect, and this inhibition of SOCE was found to be PKC-dependent (Wilson *et al*., 2015). Thus, consistent with our hypothesis for the tonic inhibition of IP_3_ formation in hepatocytes from mice fed a HFD, increased PKC activity due to elevated DAG may affect the activity of other proteins involved Ca^2+^ homeostasis and signalling. Knockout of STIM1 (the major regulator of SOCE) in lean mice was shown to impair glucose tolerance, promote insulin resistance and lead to lipid droplet accumulation, whereas STIM1 overexpression in genetically obese or HFD-fed mice ameliorated these metabolic disturbances (Arruda *et al*., 2017). These studies demonstrate that impaired Ca^2+^ homeostasis is associated with steatosis and metabolic syndrome. Whereas most previous studies have looked at basal Ca^2+^ homeostasis and alterations in Ca^2+^ signalling components in the absence of GPCR-linked agonists, our studies have directly addressed the alterations in NE-stimulated signalling following HFD. Although [Ca^2+^]_c_ oscillations depend on SERCA and SOCE to maintain and refill the ER Ca^2+^ stores, we did not observe any alterations in ER Ca^2+^ load or SOCE after short-term (1 week) HFD feeding of mice in our study (Fig. 6). Thus, the effects of HFD on IP_3_ generation appears to be an early event that precedes the changes in SERCA and SOCE, which typically occur at later times and in genetically obese animals.

We also did not detect any changes in IP_3_R density or in the amount of Ca^2+^ released from the ER with direct application of IP_3_ after short-term HFD feeding (Fig. 7), again suggesting that the reduction in NE-stimulated [Ca^2+^]_c_ oscillations is an early event in the development of NAFLD. Nevertheless, a number of studies have shown that obesity and fatty liver are associated with increased hepatic IP_3_R1 expression (Wang *et al*., 2012; Arruda *et al*., 2014; Feriod *et al*., 2017). The increased IP_3_R1 expression was shown to be associated with the ER-mitochondrial associated membranes (MAMs), which leads to enhanced Ca^2+^ transfer from ER to mitochondria and an increase in ROS production (Arruda *et al*., 2014; Feriod *et al*., 2017). These changes in IP_3_R density and MAMs may reflect an adaptive response to overcome the reduced agonist-dependent Ca^2+^ signalling described in the current study. Thus, upregulating IP_3_R1 and its localization in the MAM would help to maintain Ca^2+^-dependent stimulation of mitochondrial metabolism and oxidative phosphorylation to generate ATP. A recent longitudinal study using a high-fat plus high-sucrose diet actually reported reduce IP_3_R and MAM density at early times, and only found increased Ca^2+^ transfer to the mitochondria after 16 weeks (Beaulant *et al*., 2022).

An important question is how does a decrease in Ca^2+^ signalling capacity promote steatosis? NE and other Ca^2+^-dependent hormones such as vasopressin have been shown to inhibit acetyl-CoA-carboxylase in hepatocytes (Ma & Hems, 1975; Ly & Kim, 1981), and the net result of this inhibition is a suppression of *de novo* lipogenesis and the stimulation of β-oxidation. Thus, the loss of [Ca^2+^]_c_ signals in response to HFD could eliminate this important brake on lipogenic pathways. Consistent with this, inhibition of GPCR-dependent Ca^2+^ release leads to lipid accumulation. In *Drosophila*, IP_3_R loss of function mutants resulted in excess lipid storage and obesity (Subramanian *et al*., 2013). In mice, adipocyte-specific IP_3_R1 & 2 knockout resulted in increased weight gain after HFD due to fat accumulation (Guney *et al*., 2021). Finally, systemic glucagon infusion was shown to protect against diet-induced hepatic steatosis, an affect lost in IP_3_R1 knockout mice (Perry *et al*., 2020). There is important zonation of hepatic metabolism, such that the periportal region specializes in gluconeogenesis and β-oxidation, whereas the pericentral region specializes in lipogenesis and glycolysis (Jungermann, 1987; Berndt *et al*., 2021). It is well established that hepatic lipid accumulation first manifests in the pericentral zone in both patients and rodent models (Brunt *et al*., 1999; Adams *et al*., 2005; Chalasani *et al*., 2008; Brunt, 2010). We hypothesize that reduced propagation of lobular Ca^2+^ waves to the pericentral zone in livers of HFD-fed mice leads to a loss of catabolic regulation of lipid metabolism and excess lipid droplet accumulation specifically in this zone.

In summary, our data demonstrate that short-term HFD feeding of mice impairs NE-stimulated Ca^2+^ signalling, resulting in a 10-fold shift in EC_50_ (from 3nM to 30 nM) and suppression of robust [Ca^2+^]_c_ oscillations in isolated hepatocytes, which was also reflected in disruption of intralobular [Ca^2+^]_c_ waves in intact liver. Changes in SERCA activity, SOCE and IP_3_R density that have been associated with longer term HFD studies and genetic models of obesity were not observed with short-term HFD. Instead, we identified a novel effect of HFD to inhibit GPCR-dependent PLC activation, which resulted in significantly reduced IP_3_ generation. This suppression of NE-induced IP_3_ formation can explain the reduced [Ca^2+^]_c_ oscillation frequency in isolated hepatocytes, independent of changes in other Ca^2+^ signalling components. Moreover, in intact perfused liver, the inhibition of NE-stimulated Ca^2+^ signalling was associated with suppressed propagation of trans-lobular intercellular [Ca^2+^]_c_ waves, which may lead to dysregulation of the coordinate lobular regulation of metabolism. It is tempting to speculate that later changes in components of hepatic Ca^2+^ homeostasis, including IP_3_R upregulation, MAM formation and changes in SERCA/SOCE, reflect an adaptive response to the initial inhibition of GPCR-dependent Ca^2+^ signalling. Thus, therapeutics that rescue the upstream hormone-stimulated Ca^2+^ signalling pathways have the potential to prevent subsequent maladaptive changes to the Ca^2+^ signalling machinery and prevent or delay the onset of NAFLD.

## Data Availability Statement

All data are included within the figures and the supplementary material.

## Author contributions

R.B, A.P.T and P.J.B designed research; R.B, J.C.V, S.T, O.S and P.J.B performed research; R.B, J.C.V, S.T and P.J.B analyzed data; R.B, A.P.T and P.J.B wrote the manuscript; All authors have read and approved the final version of the manuscript.

## Acknowledgements

pGP-CMV-GCaMP6f was a gift from Douglas Kim & GENIE Project (Addgene plasmid # 40755).

## Supplementary material

Videos S1-S4 can be found here https://drive.google.com/drive/folders/1-1CGcQB5akYpFq-zPVVb2iYFA5Zuefvm?usp=sharing

**Videos S1 and S2. Image time-series showing norepinephrine-stimulated Ca^2+^ responses in perfused livers of LFD and HFD fed mice.** Livers from GCaMP6f^hep/cre^ mice fed LFD (**S1**) or HFD (**S2**) for 1 week were perfused with the indicated concentrations of NE. GCaMP6f fluorescence images were collected at 0.25 Hz using a confocal microscope focused 50-100 µm into the tissue to acquire a [Ca^2+^]_c_ image time-series, with fluorescence intensity shown in greyscale. Individual hepatocytes are easily resolved, and these are organized in multicellular cords separated by the darker sinusoidal spaces. Each of the videos covers at least two hepatic lobules (see Fig. 4A-B). As the NE concentration in the perfusion buffer is stepped up, [Ca^2+^]_c_ increases can be seen to propagate from hepatocyte to hepatocyte along the hepatic cords. [Ca^2+^]_c_ responses in individual cells from these videos are shown in Fig. 4E-F.

**Videos S3 and S4. Differential image time-series showing rate of change of Ca^2+^ in response to norepinephrine stimulation in perfused livers of LFD and HFD fed mice.** The imaging data underlying Videos S1 and S2 were processed using a differentiation algorithm to visualize the spatiotemporal organization of [Ca^2+^]_c_ increases to generate Videos S3 (LFD) and S4 (HFD), respectively. Red intensity is proportional to the rate of rise of GCaMP6f fluorescence and is overlaid on the initial pre-stimulation greyscale image to identify the location of the [Ca^2+^]_c_ increases. Note that only the rising phase of each [Ca^2+^]_c_ increase is visualized in the red differential image, so this does not show the full duration of [Ca^2+^]_c_ elevation, which can be seen in the greyscale images of Videos S1 and S2. Individual images of the initial response to 30 nM NE from these videos, and maximum intensity projections through the entire period (380 s) of exposure to 30 nM NE are shown in Fig. 4A-B.

## References

1. Adams LA, Angulo P & Lindor KD. (2005). Nonalcoholic fatty liver disease. CMAJ 172, 899–905.

2. Alcántara-Hernández R, Vázquez-Prado J & García-Sáinz JA. (2000). Protein phosphatase-protein kinase interplay modulates α_1B_-adrenoceptor phosphorylation: effects of okadaic acid. British journal of pharmacology 129, 724–730.

3. Arruda AP, Pers BM, Parlakgul G, Güney E, Goh T, Cagampan E, Lee GY, Goncalves RL & Hotamisligil GS. (2017). Defective STIM-mediated store operated Ca(2+) entry in hepatocytes leads to metabolic dysfunction in obesity. eLife 6, e29968.

4. Arruda AP, Pers BM, Parlakgul G, Guney E, Inouye K & Hotamisligil GS. (2014). Chronic enrichment of hepatic endoplasmic reticulum-mitochondria contact leads to mitochondrial dysfunction in obesity. Nat Med 20, 1427–1435.

5. Bartlett PJ, Antony AN, Agarwal A, Hilly M, Prince VL, Combettes L, Hoek JB & Gaspers LD. (2017). Chronic alcohol feeding potentiates hormone-induced calcium signalling in hepatocytes. J Physiol 595, 3143–3164.

6. Bartlett PJ, Cloete I, Sneyd J & Thomas AP. (2020). IP(3)-Dependent Ca(2+) Oscillations Switch into a Dual Oscillator Mechanism in the Presence of PLC-Linked Hormones. iScience 23, 101062.

7. Bartlett PJ, Gaspers LD, Pierobon N & Thomas AP. (2014). Calcium-dependent regulation of glucose homeostasis in the liver. Cell Calcium 55, 306–316.

8. Bartlett PJ, Metzger W, Gaspers LD & Thomas AP. (2015). Differential regulation of multiple steps in inositol 1,4,5-trisphosphate signaling by protein kinase C shapes hormone-stimulated Ca2+ oscillations. Journal of Biological Chemistry.

9. Bartlett PJ, Young KW, Nahorski SR & Challiss RA. (2005). Single cell analysis and temporal profiling of agonist-mediated inositol 1,4,5-trisphosphate, Ca2+, diacylglycerol, and protein kinase C signaling using fluorescent biosensors. J Biol Chem 280, 21837-21846.

10. Beaulant A, Dia M, Pillot B, Chauvin M-A, Ji-cao J, Durand C, Bendridi N, Chanon S, Vieille-Marchiset A, Da Silva CC, Patouraux S, Anty R, Iannelli A, Tran A, Gual P, Vidal H, Gomez L, Paillard M & Rieusset J. (2022). Endoplasmic reticulum-mitochondria miscommunication is an early and causal trigger of hepatic insulin resistance and steatosis. J Hepatol.

11. Ben-Moshe S, Shapira Y, Moor AE, Manco R, Veg T, Bahar Halpern K & Itzkovitz S. (2019). Spatial sorting enables comprehensive characterization of liver zonation. Nature Metabolism 1, 899–911.

12. Berndt N, Kolbe E, Gajowski R, Eckstein J, Ott F, Meierhofer D, Holzhütter H-G & Matz-Soja M. (2021). Functional Consequences of Metabolic Zonation in Murine Livers: Insights for an Old Story. Hepatology 73, 795–810.

13. Berrie CP & Cobbold PH. (1995). Both activators and inhibitors of protein kinase C promote the inhibition of phenylephrine-induced [Ca2+]i oscillations in single intact rat hepatocytes. Cell Calcium 18, 232–244.

14. Betzenhauser MJ, Fike JL, Wagner LE & Yule DI. (2009). Protein Kinase A Increases Type-2 Inositol 1,4,5-Trisphosphate Receptor Activity by Phosphorylation of Serine 937. Journal of Biological Chemistry 284, 25116-25125.

15. Bezprozvanny l, Watras J & Ehrlich BE. (1991). Bell-shaped calcium-response curves of lns(l,4,5)P3- and calcium-gated channels from endoplasmic reticulum of cerebellum. Nature 351, 751-754.

16. Blackmore PF, Strickland WG, Bocckino SB & Exton JH. (1986). Mechanism of hepatic glycogen synthase inactivation induced by Ca2+-mobilizing hormones. Studies using phospholipase C and phorbol myristate acetate. Biochem J 237, 235–242.

17. Boden G, She P, Mozzoli M, Cheung P, Gumireddy K, Reddy P, Xiang X, Luo Z & Ruderman N. (2005). Free Fatty Acids Produce Insulin Resistance and Activate the Proinflammatory Nuclear Factor-κB Pathway in Rat Liver. Diabetes 54, 3458–3465.

18. Bredt DS, Mourey RJ & Snyder SH. (1989). A simple, sensitive, and specific radioreceptor assay for inositol 1,4,5-trisphosphate in biological tissues. Biochemical and Biophysical Research Communications 159, 976-982.

19. Brownsey RW, Boone AN, Elliott JE, Kulpa JE & Lee WM. (2006). Regulation of acetyl-CoA carboxylase. Biochemical Society transactions 34, 223–227.

20. Brunt EM. (2010). Pathology of nonalcoholic fatty liver disease. Nat Rev Gastroenterol Hepatol 7.

21. Brunt EM, Janney CG, Di Bisceglie AM, Neuschwander-Tetri BA & Bacon BR. (1999). Nonalcoholic steatohepatitis: a proposal for grading and staging the histological lesions. Am J Gastroenterol 94.

22. Cardenas C, Miller RA, Smith I, Bui T, Molgo J, Muller M, Vais H, Cheung KH, Yang J, Parker I, Thompson CB, Birnbaum MJ, Hallows KR & Foskett JK. (2010). Essential regulation of cell bioenergetics by constitutive InsP3 receptor Ca2+ transfer to mitochondria. Cell 142, 270–283.

23. Chalasani N, Wilson L, Kleiner DE, Cummings OW, Brunt EM, Unalp A & Network NCR. (2008). Relationship of steatosis grade and zonal location to histological features of steatohepatitis in adult patients with non-alcoholic fatty liver disease. J Hepatol 48, 829–834.

24. Chen TW, Wardill TJ, Sun Y, Pulver SR, Renninger SL, Baohan A, Schreiter ER, Kerr RA, Orger MB, Jayaraman V, Looger LL, Svoboda K & Kim DS. (2013). Ultrasensitive fluorescent proteins for imaging neuronal activity. Nature 499, 295–300.

25. Cloete I, Bartlett PJ, Kirk V, Thomas AP & Sneyd J. (2020). Dual mechanisms of Ca2+ oscillations in hepatocytes. Journal of Theoretical Biology 503, 110390.

26. Cloete I, Corrêa-Velloso JC, Bartlett PJ, Kirk V, Thomas AP & Sneyd J. (2021). A Tale of two receptors. J Theor Biol 518, 110629.

27. Corrêa-Velloso JC, Bartlett PJ, Brumer R, Gaspers LD, Ulrich H & Thomas AP. (2021). Receptor-specific Ca(2+) oscillation patterns mediated by differential regulation of P2Y purinergic receptors in rat hepatocytes. iScience 24, 103139.

28. Dai W, Ye L, Liu A, Wen SW, Deng J, Wu X & Lai Z. (2017). Prevalence of nonalcoholic fatty liver disease in patients with type 2 diabetes mellitus: A meta-analysis. Medicine (Baltimore) 96, e8179.

29. Denton RM. (2009). Regulation of mitochondrial dehydrogenases by calcium ions. Biochim Biophys Acta 1787, 1309–1316.

30. Fazel Y, Koenig AB, Sayiner M, Goodman ZD & Younossi ZM. (2016). Epidemiology and natural history of non-alcoholic fatty liver disease. Metabolism 65, 1017–1025.

31. Feriod CN, Gustavo Oliveira A, Guerra MT, Nguyen L, Mitchell Richards K, Jurczak MJ, Ruan H-B, Paulo Camporez J, Yang X, Shulman GI, Bennett AM, Nathanson MH & Ehrlich BE. (2017). Hepatic inositol 1,4,5 trisphosphate receptor type 1 mediates fatty liver. Hepatology Communications 1, 23-35.

32. Feriod CN, Nguyen L, Jurczak MJ, Kruglov EA, Nathanson MH, Shulman GI, Bennett AM & Ehrlich BE. (2014). Inositol 1,4,5-trisphosphate receptor type II (InsP3R-II) is reduced in obese mice, but metabolic homeostasis is preserved in mice lacking InsP3R-II. American Journal of Physiology - Endocrinology and Metabolism 307, E1057-E1064.

33. Foskett JK, White C, Cheung KH & Mak DO. (2007). Inositol trisphosphate receptor Ca2+ release channels. Physiol Rev 87, 593–658.

34. Fu S, Yang L, Li P, Hofmann O, Dicker L, Hide W, Lin X, Watkins SM, Ivanov AR & Hotamisligil GS. (2011). Aberrant lipid metabolism disrupts calcium homeostasis causing liver endoplasmic reticulum stress in obesity. Nature 473, 528–531.

35. Fung TT, Rimm EB, Spiegelman D, Rifai N, Tofler GH, Willett WC & Hu FB. (2001). Association between dietary patterns and plasma biomarkers of obesity and cardiovascular disease risk. Am J Clin Nutr 73, 61–67.

36. Gaspers LD, Bartlett PJ, Politi A, Burnett P, Metzger W, Johnston J, Joseph SK, Hofer T & Thomas AP. (2014). Hormone-induced calcium oscillations depend on cross-coupling with inositol 1,4,5-trisphosphate oscillations. Cell reports 9, 1209-1218.

37. Gaspers LD, Pierobon N & Thomas AP. (2019). Intercellular calcium waves integrate hormonal control of glucose output in the intact liver. J Physiol 597, 2867–2885.

38. Gaspers LD & Thomas AP. (2005). Calcium signaling in liver. Cell Calcium 38, 329–342.

39. Gaspers LD, Thomas AP, Hoek JB & Bartlett PJ. (2021). Ethanol disrupts hormone-induced calcium signaling in liver. Function.

40. Geraldes P & King GL. (2010). Activation of Protein Kinase C Isoforms & Its Impact on Diabetic Complications. Circ Res 106, 1319–1331.

41. Greene MW, Burrington CM, Ruhoff MS, Johnson AK, Chongkrairatanakul T & Kangwanpornsiri A. (2010). PKCδ Is Activated in a Dietary Model of Steatohepatitis and Regulates Endoplasmic Reticulum Stress and Cell Death. J Biol Chem 285, 42115–42129.

42. Guney E, Arruda AP, Parlakgul G, Cagampan E, Min N, Lee GY, Greene L, Tsaousidou E, Inouye K, Han MS, Davis RJ & Hotamisligil GS. (2021). Aberrant Ca^2+^ signaling by IP 3Rs in adipocytes links inflammation to metabolic dysregulation in obesity. Sci Signal 14, eabf2059.

43. Hajnóczky G, Gao E, Nomura T, Hoek JB & Thomas AP. (1993). Multiple mechanisms by which protein kinase A potentiates inositol 1,4,5-trisphosphate-induced Ca^2+^ mobilization in permeabilized hepatocytes. Biochemical Journal 293, 413-422.

44. Hajnóczky G, Lin C & Thomas AP. (1994). Luminal communication between intracellular calcium stores modulated by GTP and the cytoskeleton. Journal of Biological Chemistry 269, 10280–10287.

45. Hajnoczky G, Robb-Gaspers LD, Seitz MB & Thomas AP. (1995). Decoding of cytosolic calcium oscillations in the mitochondria. Cell 82, 415–424.

46. Hajnóczky G & Thomas AP. (1997). Minimal requirements for calcium oscillations driven by the IP_3_ receptor. Embo J 16, 3533–3543.

47. Institute of Medicine (U.S.). Panel on Macronutrients. & Institute of Medicine (U.S.). Standing Committee on the Scientific Evaluation of Dietary Reference Intakes. (2005). Dietary reference intakes for energy, carbohydrate, fiber, fat, fatty acids, cholesterol, protein, and amino acids. National Academies Press, Washington, D.C.

48. Itani SI, Ruderman NB, Schmieder F & Boden G. (2002). Lipid-Induced Insulin Resistance in Human Muscle Is Associated With Changes in Diacylglycerol, Protein Kinase C, and IκB-α. Diabetes 51, 2005–2011.

49. Jouaville LS, Pinton P, Bastianutto C, Rutter GA & Rizzuto R. (1999). Regulation of mitochondrial ATP synthesis by calcium: Evidence for a long-term metabolic priming. Proceedings of the National Academy of Sciences 96, 13807–13812.

50. Jungermann K. (1987). Metabolic zonation of liver parenchyma: Significance for the regulation of glycogen metabolism, gluconeogenesis, and glycolysis. Diabetes/Metabolism Reviews 3, 269–293.

51. Khamphaya T, Chukijrungroat N, Saengsirisuwan V, Mitchell-Richards KA, Robert ME, Mennone A, Ananthanarayanan M, Nathanson MH & Weerachayaphorn J. (2018). Nonalcoholic fatty liver disease impairs expression of the type II inositol 1,4,5-trisphosphate receptor. Hepatology 67, 560-574.

52. Khan MT, Wagner L, Yule DI, Bhanumathy C & Joseph SK. (2006). Akt Kinase Phosphorylation of Inositol 1,4,5-Trisphosphate Receptors. Journal of Biological Chemistry 281, 3731-3737.

53. Kraus-Friedmann N & Feng L. (1996). The role of intracellular Ca2+ in the regulation of gluconeogenesis. Metabolism 45, 389–403.

54. Kumashiro N, Erion DM, Zhang D, Kahn M, Beddow SA, Chu X, Still CD, Gerhard GS, Han X, Dziura J, Petersen KF, Samuel VT & Shulman GI. (2011). Cellular mechanism of insulin resistance in nonalcoholic fatty liver disease. Proceedings of the National Academy of Sciences 108, 16381–16385.

55. Li W-h, Llopis J, Whitney M, Zlokarnik G & Tsien RY. (1998). Cell-permeant caged InsP3 ester shows that Ca2+ spike frequency can optimize gene expression. Nature 392, 936–941.

56. Lu Q, Tian X, Wu H, Huang J, Li M, Mei Z, Zhou L, Xie H & Zheng S. (2021). Metabolic Changes of Hepatocytes in NAFLD. Frontiers in Physiology 12.

57. Ly S & Kim KH. (1981). Inactivation of hepatic acetyl-CoA carboxylase by catecholamine and its agonists through the alpha-adrenergic receptors. Journal of Biological Chemistry 256, 11585–11590.

58. Ma GY & Hems DA. (1975). Inhibition of fatty acid synthesis and stimulation of glycogen breakdown by vasopressin in the perfused mouse liver. Biochemical Journal 152, 389–392.

59. Miller RA & Birnbaum MJ. (2016). Glucagon: acute actions on hepatic metabolism. Diabetologia 59, 1376–1381.

60. Muzurović E, Mikhailidis DP & Mantzoros C. (2021). Non-Alcoholic Fatty Liver Disease, Insulin Resistance, Metabolic Syndrome and their Association with Vascular Risk. Metabolism, 154770.

61. Ozcan L, Wong Catherine CL, Li G, Xu T, Pajvani U, Park Sung Kyu R, Wronska A, Chen B-X, Marks Andrew R, Fukamizu A, Backs J, Singer Harold A, Yates Iii John R, Accili D & Tabas I. (2012). Calcium Signaling through CaMKII Regulates Hepatic Glucose Production in Fasting and Obesity. Cell Metabolism 15, 739-751.

62. Park SW, Zhou Y, Lee J, Lee J & Ozcan U. (2010). Sarco(endo)plasmic reticulum Ca2+-ATPase 2b is a major regulator of endoplasmic reticulum stress and glucose homeostasis in obesity. Proc Natl Acad Sci U S A 107, 19320–19325.

63. Perry RJ, Zhang D, Guerra MT, Brill AL, Goedeke L, Nasiri AR, Rabin-Court A, Wang Y, Peng L, Dufour S, Zhang Y, Zhang XM, Butrico GM, Toussaint K, Nozaki Y, Cline GW, Petersen KF, Nathanson MH, Ehrlich BE & Shulman GI. (2020). Glucagon stimulates gluconeogenesis by INSP3R1-mediated hepatic lipolysis. Nature 579, 279–283.

64. Politi A, Gaspers LD, Thomas AP & Hofer T. (2006). Models of IP_3_ and Ca2+ oscillations: frequency encoding and identification of underlying feedbacks. Biophys J 90, 3120–3133.

65. Ritter O, Jelenik T & Roden M. (2015). Lipid-mediated muscle insulin resistance: different fat, different pathways? Journal of Molecular Medicine 93, 831–843.

66. Robb-Gaspers LD, Burnett P, Rutter GA, Denton RM, Rizzuto R & Thomas AP. (1998a). Integrating cytosolic calcium signals into mitochondrial metabolic responses. Embo J 17, 4987–5000.

67. Robb-Gaspers LD, Rutter GA, Burnett P, Hajnoczky G, Denton RM & Thomas AP. (1998b). Coupling between cytosolic and mitochondrial calcium oscillations: role in the regulation of hepatic metabolism. Biochim Biophys Acta 1366, 17–32.

68. Robb-Gaspers LD & Thomas AP. (1995). Coordination of Ca2+ signaling by intercellular propagation of Ca2+ waves in the intact liver. J Biol Chem 270, 8102–8107.

69. Rooney TA, Joseph SK, Queen C & Thomas AP. (1996). Cyclic GMP Induces Oscillatory Calcium Signals in Rat Hepatocytes. Journal of Biological Chemistry 271, 19817–19825.

70. Rooney TA, Sass EJ & Thomas AP. (1989). Characterization of cytosolic calcium oscillations induced by phenylephrine and vasopressin in single fura-2-loaded hepatocytes. J Biol Chem 264, 17131–17141.

71. Rui L. (2014). Energy Metabolism in the Liver. Comprehensive Physiology 4, 177–197.

72. Samuel VT, Liu Z-X, Wang A, Beddow SA, Geisler JG, Kahn M, Zhang X-m, Monia BP, Bhanot S & Shulman GI. (2007). Inhibition of protein kinase Cε prevents hepatic insulin resistance in nonalcoholic fatty liver disease. The Journal of Clinical Investigation 117, 739–745.

73. Sanchez-Bueno A, Dixon CJ, Woods NM, Cuthbertson KS & Cobbold PH. (1990). Inhibitors of protein kinase C prolong the falling phase of each free-calcium transient in a hormone-stimulated hepatocyte. Biochem J 268, 627–632.

74. Sayiner M, Koenig A, Henry L & Younossi ZM. (2016). Epidemiology of Nonalcoholic Fatty Liver Disease and Nonalcoholic Steatohepatitis in the United States and the Rest of the World. Clin Liver Dis 20, 205–214.

75. Schmitz-Peiffer C & Biden TJ. (2010). PKCδ Blues for the β-Cell. Diabetes 59, 1–3.

76. Sharara-Chami RI, Joachim M, Pacak K & Majzoub JA. (2010). Glucocorticoid treatment--effect on adrenal medullary catecholamine production. Shock 33, 213–217.

77. Strassheim D & Williams CL. (2000). P2Y2 purinergic and M3 muscarinic acetylcholine receptors activate different phospholipase C-beta isoforms that are uniquely susceptible to protein kinase C-dependent phosphorylation and inactivation. J Biol Chem 275, 39767–39772.

78. Subramanian M, Metya SK, Sadaf S, Kumar S, Schwudke D & Hasan G. (2013). Altered lipid homeostasis in Drosophila InsP3 receptor mutants leads to obesity and hyperphagia. Disease Models & Mechanisms 6, 734–744.

79. Thomas AP, Renard-Rooney DC, Hajnoczky G, Robb-Gaspers LD, Lin C & Rooney TA. (1995). Subcellular organization of calcium signalling in hepatocytes and the intact liver. Ciba Foundation symposium 188, 18–35; discussion 35-49.

80. Thomas AP, Renard DC & Rooney TA. (1991). Spatial and temporal organization of calcium signalling in hepatocytes. Cell Calcium 12, 111–126.

81. Thurley K, Tovey SC, Moenke G, Prince VL, Meena A, Thomas AP, Skupin A, Taylor CW & Falcke M. (2014). Reliable encoding of stimulus intensities within random sequences of intracellular Ca2+ spikes. Sci Signal 7, ra59.

82. Turban S & Hajduch E. (2011). Protein kinase C isoforms: Mediators of reactive lipid metabolites in the development of insulin resistance. FEBS Lett 585, 269–274.

83. van Schadewijk A, van’t Wout EFA, Stolk J & Hiemstra PS. (2012). A quantitative method for detection of spliced X-box binding protein-1 (XBP1) mRNA as a measure of endoplasmic reticulum (ER) stress. Cell Stress & Chaperones 17, 275–279.

84. Vandesompele J, De Preter K, Pattyn F, Poppe B, Van Roy N, De Paepe A & Speleman F. (2002). Accurate normalization of real-time quantitative RT-PCR data by geometric averaging of multiple internal control genes. Genome Biol 3, Research0034.

85. Vollmers C, Gill S, DiTacchio L, Pulivarthy SR, Le HD & Panda S. (2009). Time of feeding and the intrinsic circadian clock drive rhythms in hepatic gene expression. Proc Natl Acad Sci U S A 106, 21453–21458.

86. Wang Y, Li G, Goode J, Paz JC, Ouyang K, Screaton R, Fischer WH, Chen J, Tabas I & Montminy M. (2012). Inositol-1,4,5-trisphosphate receptor regulates hepatic gluconeogenesis in fasting and diabetes. Nature 485, 128-132.

87. Wilson Claire H, Ali Eunüs S, Scrimgeour N, Martin Alyce M, Hua J, Tallis George A, Rychkov Grigori Y & Barritt Greg J. (2015). Steatosis inhibits liver cell store-operated Ca2+ entry and reduces ER Ca2+ through a protein kinase C-dependent mechanism. Biochemical Journal 466, 379–390.

88. Wong RJ, Aguilar M, Cheung R, Perumpail RB, Harrison SA, Younossi ZM & Ahmed A. (2015). Nonalcoholic steatohepatitis is the second leading etiology of liver disease among adults awaiting liver transplantation in the United States. Gastroenterology 148, 547–555.

89. Young KW, Nash MS, Challiss RA & Nahorski SR. (2003). Role of Ca2+ feedback on single cell inositol 1,4,5-trisphosphate oscillations mediated by G-protein-coupled receptors. J Biol Chem 278, 20753-20760.

90. Yu C, Chen Y, Cline GW, Zhang D, Zong H, Wang Y, Bergeron R, Kim JK, Cushman SW, Cooney GJ, Atcheson B, White MF, Kraegen EW & Shulman GI. (2002). Mechanism by Which Fatty Acids Inhibit Insulin Activation of Insulin Receptor Substrate-1 (IRS-1)-associated Phosphatidylinositol 3-Kinase Activity in Muscle. Journal of Biological Chemistry 277, 50230–50236.

91. Zhang J, Li Y, Jiang S, Yu H & An W. (2014). Enhanced endoplasmic reticulum SERCA activity by overexpression of hepatic stimulator substance gene prevents hepatic cells from ER stress-induced apoptosis. American Journal of Physiology-Cell Physiology 306, C279–C290.

